# Tumor suppressor heterozygosity and homologous recombination deficiency mediate resistance to front-line therapy in breast cancer

**DOI:** 10.1101/2024.02.05.578934

**Authors:** Anton Safonov, Antonio Marra, Chaitanya Bandlamudi, Ben O’Leary, Bradley Wubbenhorst, Emanuela Ferraro, Enrico Moiso, Minna Lee, Julia An, Mark T.A. Donoghue, Marie Will, Fresia Pareja, Emily Nizialek, Natalia Lukashchuk, Eleni Sofianopoulou, Yuan Liu, Xin Huang, Mehnaj Ahmed, Miika M. Mehine, Dara Ross, Diana Mandelker, Marc Ladanyi, Nikolaus Schultz, Michael F. Berger, Maurizio Scaltriti, Jorge S. Reis-Filho, Bob T. Li, Ken Offit, Larry Norton, Ronglai Shen, Sohrab Shah, Kara N. Maxwell, Fergus Couch, Susan M. Domchek, David B. Solit, Katherine L. Nathanson, Mark E. Robson, Nicholas C. Turner, Sarat Chandarlapaty, Pedram Razavi

## Abstract

The co-occurrence of germline and somatic oncogenic alterations is frequently observed in breast cancer, but their combined biologic and clinical significance has not been evaluated. To assess the role of germline-somatic interactions on outcomes in routine practice, we developed an integrated clinicogenomic pipeline to analyze the genomes of over 4,500 patients with breast cancer. We find that germline (g)*BRCA2*-associated tumors are enriched for *RB1* loss-of-function mutations and manifest poor outcomes on standard-of-care, front-line CDK4/6 inhibitor (CDK4/6i) combinations. Amongst these tumors, g*BRCA2*-related homologous recombination deficiency (HRD) as well as baseline *RB1* LOH status promote acquisition of *RB1* loss-of-function mutations under the selective pressure of CDK4/6i, causing therapy resistance. These findings suggest an alternative therapeutic strategy using sequential targeting of HRD in g*BRCA-* associated breast cancers through PARP inhibitors *prior to* CDK4/6i therapy to intercept deleterious *RB1*-loss trajectories and thus suppress the emergence of CDK4/6 inhibitor resistance. More broadly, our findings demonstrate how germline-somatic driven genomic configurations shape response to systemic therapy and can be exploited therapeutically as part of biomarker-directed clinical strategies.

## INTRODUCTION

Inherited pathogenic variants (PVs) in cancer susceptibility genes predispose to tumor development^1,2^. Cancer susceptibility genes often play fundamental roles in DNA damage repair^3–6^ and regulation of cell cycle progression^7–9^. In parallel, *somatic* alterations affecting many of the same DNA and cell cycle regulatory processes are observed in most fully established cancers. Recent studies have begun to study the interplay between germline and somatic features, exploring how the landscape of somatic variants is shaped by a patient’s germline genetic profile^10–15^. Yet, these broad pan-cancer analyses have not fully considered biologically and clinically pertinent cancer-type-specific tumor characteristics, such as receptor subtypes in breast cancer. Furthermore, the therapeutic ramifications of germline-somatic interactions remain underexplored, even though both somatic and germline variants are now recognized as predictive biomarkers of treatment response. Targeted therapies are now available to exploit therapeutic vulnerabilities stemming from pathogenic *germline* alterations, such as PARP inhibitors (PARPi) for the treatment of homologous recombination (HRD)-driven tumors arising in patients with germline *BRCA2* mutation (*gBRCA2*)^16,17^. Conversely, somatic variants not only serve as actionable targets themselves, but also can emerge as mechanisms of resistance to targeted therapies^18^.

An abundance of choices between approved targeted and lineage-directed therapies can lead to ambiguity in selecting the optimal sequencing of therapies. This clinical dilemma is exemplified in the management of g*BRCA2*-driven breast cancers, the majority of which have hormone receptor overexpression (HR+) and are thus sensitive to endocrine therapies (ET)^19^. The combination of CDK4/6 inhibitors (CDK4/6i) and ET is a standard-of-care front-line treatment strategy for breast cancers in both the metastatic and adjuvant treatment setting^20–22^; however, it is unclear whether this strategy should be deployed prior to or in lieu of PARPi, which are also approved for patients with ER+ breast cancers arising in the setting of *gBRCA2* mutation.

Specifically, it has yet to be systematically evaluated whether the somatic variants associated with g*BRCA2* condition response or resistance to standard-of-care front line systemic therapies.

On a more general level, defining the role of pre-treatment germline and somatic genomic context on evolutionary trajectories of *acquired* treatment resistance could allow for early interception strategies designed to prevent or delay the emergence of drug resistant tumor clones. To elucidate the therapeutic relevance of the underlying germline process to the somatic genotype in a clinically relevant context, we performed an integrated analysis of germline and somatic next generation sequencing data, paired with detailed clinical data including treatment response data.

## MAIN

### Clinicopathological and genomic features of germline HRD-related variants

To identify the interactions between germline and somatic pathogenic variants in breast cancer, we integrated detailed clinical annotation with prospectively collected sequencing data from 4,640 breast cancer patients (the MSK breast cancer cohort, **Fig S1)**. DNA derived from tumor tissue and blood as a source of germline DNA were sequenced using an FDA authorized clinical sequencing assay encompassing up to 468 cancer-associated genes, including germline analysis of 84 established cancer predisposition genes^23–25^. Genes of interest for the germline analysis included canonical members of the homologous recombination pathway: g*BRCA2* (pathogenic variants in 127 patients; 2.7% of cohort), g*BRCA1* (n = 105, 2.2%), g*CHEK2* (n = 97, 2.1%), g*ATM* (n = 45, 1.0%), and g*PALB2* (n = 25, 0.5%).

The clinicopathologic characteristics of the germline-altered cancers and germline wild-type (gWT) cancers strongly reflected previously established patterns suggesting that our cohort was representative of broader population of patients with breast cancer (**Table 1**). Specifically, we observed a younger age of diagnosis with g*BRCA1/2* carriers compared to those gWT. g*BRCA1*- associated breast cancers tended to be triple-negative^26,27^ and high-grade invasive ductal carcinomas. Meanwhile, g*BRCA2*, g*CHEK2* and g*ATM* carriers typically had HR+/HER2- disease (71.0%, 74.6%, 75.6%, respectively), all consistent with prior studies^28,29^.

**Table 1.**
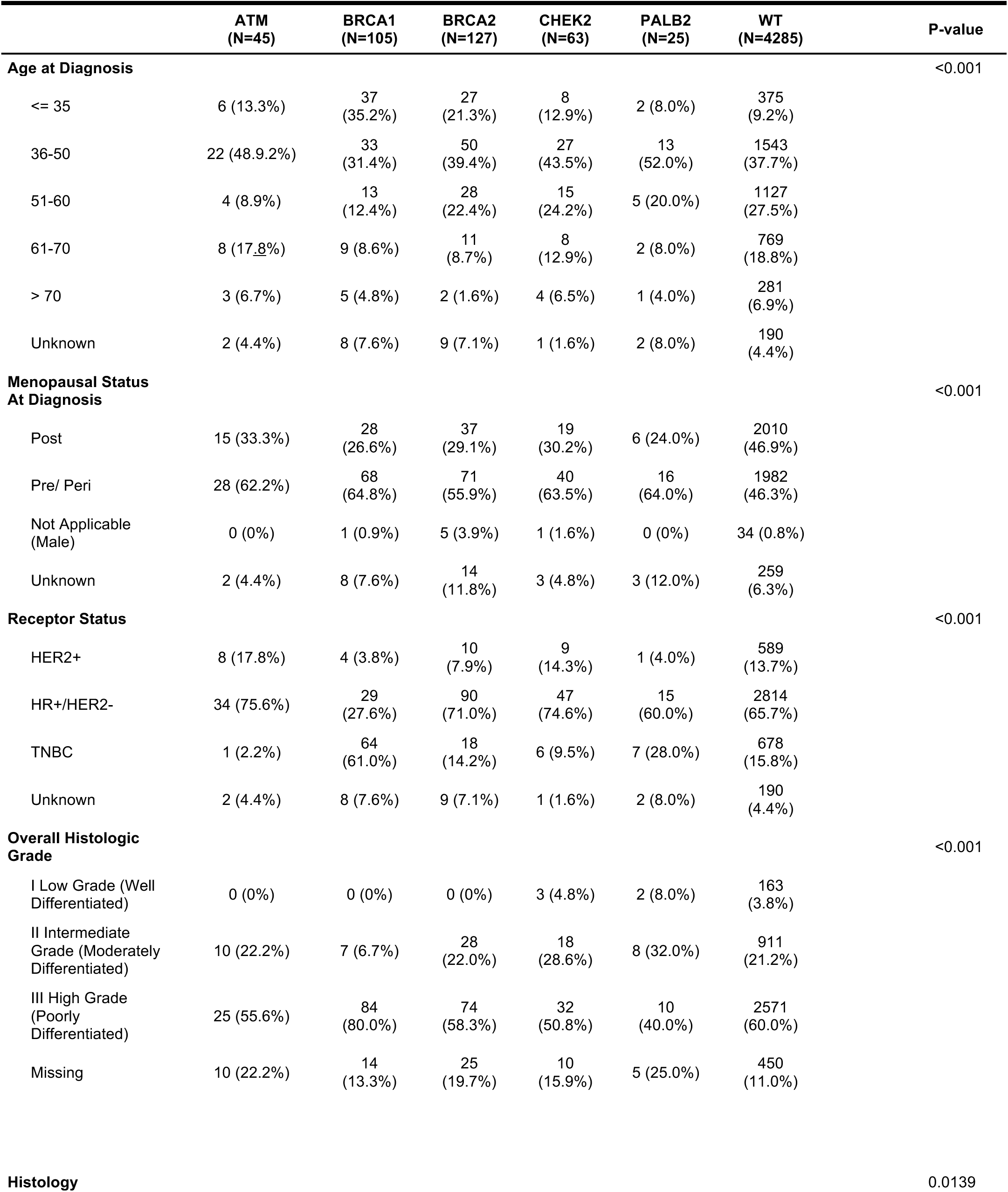
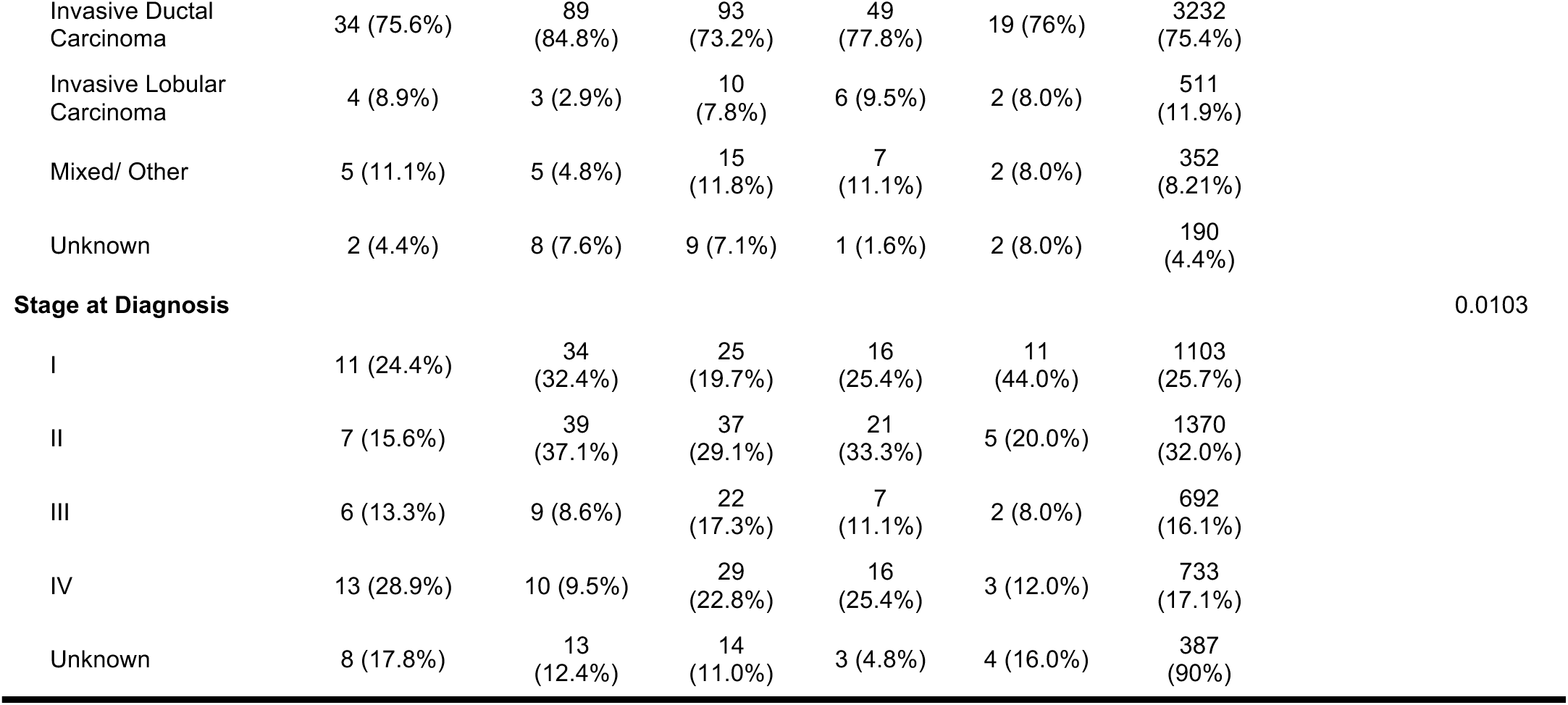

Biallelic loss is often necessary for phenotypic impact in carriers of germline pathogenic variants (PVs) in HRD-related genes, yet its incidence varies by gene, with higher rates of biallelic inactivation observed in high-penetrance genes^30,31^. Our results confirmed that biallelic inactivation rates varied significantly across genes, ranging from 47.5% in g*CHEK2* carriers, to 88.8% and 89.7% for g*BRCA1* and g*BRCA2* carriers, respectively (**Fig S2)**. We also found lower frequency of loss of heterozygosity (LOH) in g*PALB2* carriers (n = 11, 50.0%), but a relatively higher frequency of “second-hit” somatic mutations resulting in biallelic loss (n = 3, 13.6%), aligning with established patterns^32^. In confirming these previously established histologic, demographic, and genomic patterns associated with germline PVs, the clinical relevance of this clinicogenomic cohort was established.

### Analysis of germline-somatic interactions uncovers occult clinical actionability

We first sought to define patterns of mutual exclusivity or enrichment of somatic variants with germline pathogenic variants (germline-somatic interactions), in the context of receptor subtype and zygosity. We therefore compiled mutations, structural variants and copy number alterations predicted to be functionally significant based on OncoKB^33^, and performed enrichment analyses using Firth-penalized logistic regression (**Fig. 1a)**. This analysis robustly validated previously reported enrichment of *TP53* alterations in *gBRCA1* carriers and mutual exclusivity of *gATM* and *TP53* alterations^34–36^ (**Fig. 1b,c**). Both findings were more pronounced in tumors exhibiting biallelic inactivation of the respective genes^31,37^ (**Fig. S3**). We further explored these observations by conducting receptor subtype-dependent analyses of germline-somatic interactions. g*BRCA1*:*TP53* enrichment was enriched in HR+/HER2- tumors but did not meet statistical significance in in triple-negative tumors, where *TP53* is already highly prevalent in WT tumors (**Fig. S3**), supporting *TP53* loss-of-function as a fundamental event in the oncogenesis of *gBRCA1*-driven breast cancers^38–40^.

**Fig. 1:**
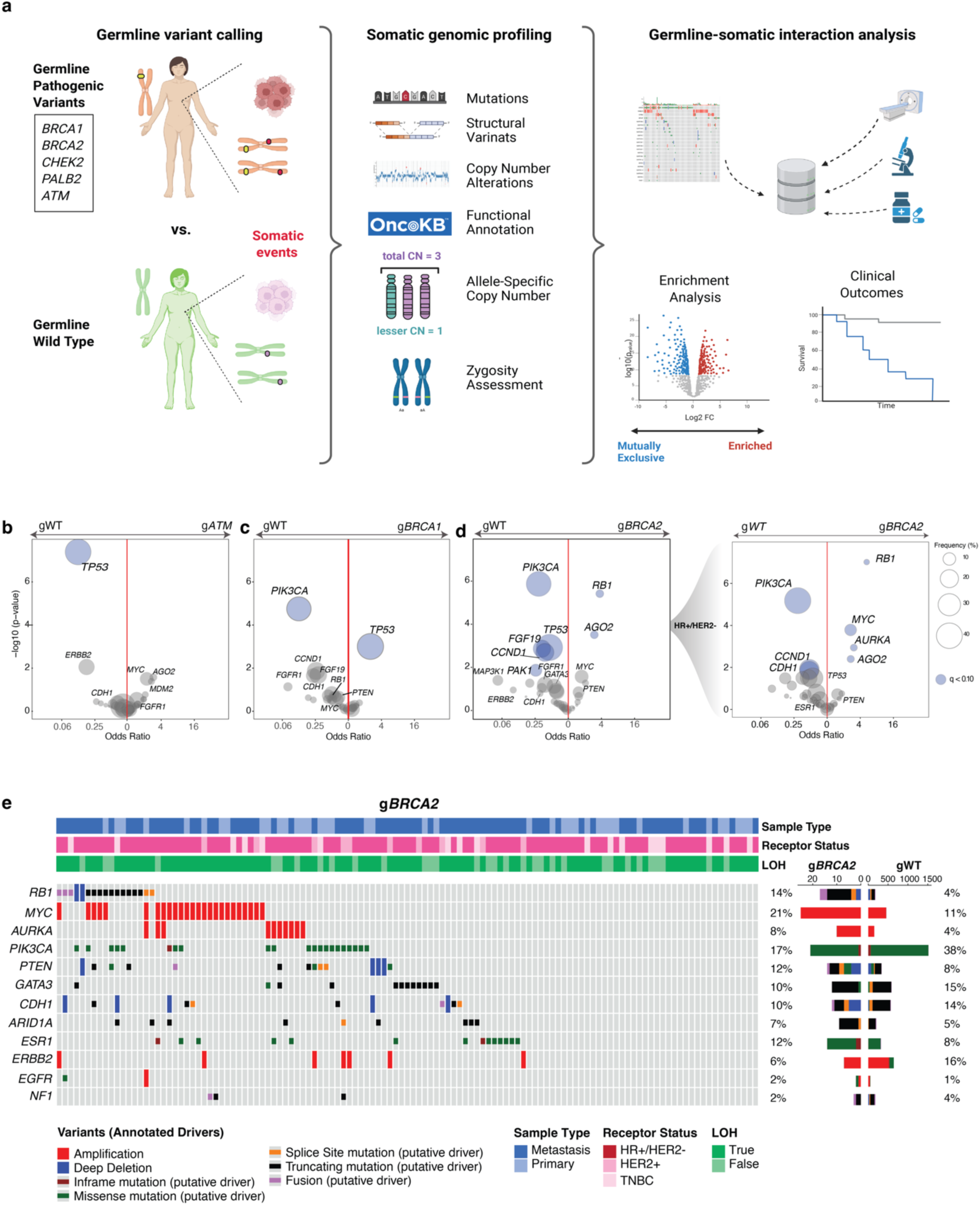
Germline-somatic interactions in breast cancer a) Study schema: Enrichment analysis was performed to identify somatic alterations more prevalent in patients with pathogenic germline variants of homologous recombination pathway (HRD) genes (compared to “sporadic” or germline wild-type cases). **b)** Results of enrichment analysis of g*BRCA1;* n = 101 compared to g*WT*; n = 4115), as determined by Firth penalized logistic regression with sample type and receptor status as covariates. Significant genes (blue) defined as q-value < 0.10. *gBRCA1* tumors were enriched for somatic *TP53* variants (2.79, (95% CI 1.59 – 4.91), q= 0.006) and mutually exclusive for *PIK3CA* variants. **c)** As in b, but depicting comparison of g*ATM,* n = 43; compared to g*WT*). **d)** As in b and c, but depicting comparison of g*BRCA2*, n = 122; compared to g*WT*, n = 4115). A subset analysis of HR+/HER2- tumors (total n = 2910, n*BRCA2* = 93, n*WT* = 2817) was then performed. Somatic *RB1* variants represented the most significant enrichment in g*BRCA2* carriers (OR 3.41 (1.99 – 5.85, q = 0.000) **e)** Oncoprint detailing mutations, copy number deletions, fusions in the indicated genes in g*BRCA2* vs. g*WT*. Receptor status, sample type, and zygosity are annotated above.

In g*BRCA2-*driven breast cancers, *RB1* somatic variants were significantly enriched, in stark contrast to their depletion in g*BRCA1*-associated tumors (**Fig. 1c,d**). Focusing on patients with HR+/HER2 tumors (71.0% of *gBRCA2*) revealed an in even more apparent enrichment of somatic *RB1* alterations in *gBRCA2* carriers. This analysis also uncovered an enrichment of *MYC* and *AURKA* amplifications in this germline context. Of clinical relevance, *RB1*, *MYC* and *AURKA* alterations have all been implicated in CDK4/6i resistance^41–43^. Conversely, *PIK3CA* alterations were enriched in the gWT cancers compared to *gBRCA2* and g*BRCA1*-driven cancers.

**Fig. 1e** displays the somatic variant type and pathologic features of g*BRCA2*-associated breast cancers. The receptor status and LOH-specific germline-somatic interactions are further displayed in **Fig. S4-S7**; corresponding Oncoprints are detailed in **Fig. S8.**

### Germline *BRCA2* driven breast cancers exhibit resistance to standard frontline CDK4/6 inhibitor-based combination therapy

*RB1* loss is an established mechanism of resistance to CDK4/6i inhibitors^45^. Based on our results demonstrating a significant enrichment of *RB1* alterations in g*BRCA2* carriers, we analyzed the impact of g*BRCA2* status on progression free survival (PFS) in patients with HR+/HER2- metastatic breast cancer (MBC) treated with CDK4/6i plus ET combinations in our cohort. A total of 1,097 breast cancer patients treated with CDK4/6i, including 533 in the first-line setting, were evaluable for breast cancer specific outcomes. g*BRCA2* PVs were associated with a significantly worse PFS on CDK4/6i and ET in univariate and multivariate analyses (median PFS 9.0 vs 13.7 months, multivariate Hazard Ratio [HR] 2.28, 95% CI 1.52 – 3.39, p = 4.9e-5) (**Fig. 2a**). Similar results were seen when the analysis was extended to all treatment lines, with consideration of ET partner and treatment line as covariates (HR 2.02 95% CI 1.52 – 2.70, p = 1.5e-6) (**Fig. 2b**). *RB1* loss-of-function mutations were rarely identified in pre-treatment samples; these results therefore held true when omitting samples where this was the case.

**Fig. 2:**
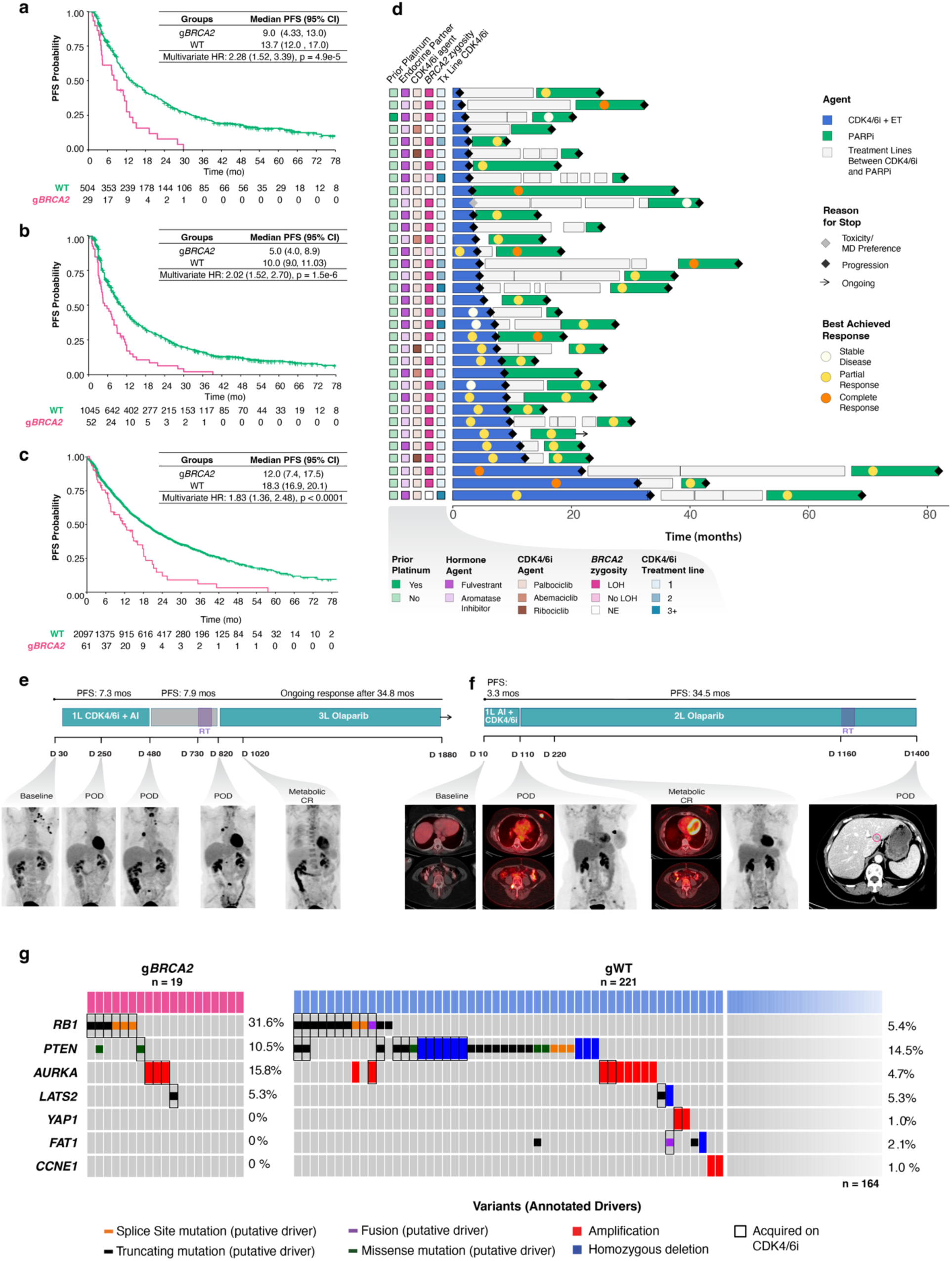
Clinical implications of g*BRCA2* status a) Progression free survival for patients treated with first-line CDK4/6i plus endocrine combination therapy by g*BRCA2* status. Multivariate Cox-proportional hazard ratio, including endocrine therapy partner as covariate (hazard ratio 2.28, 95% confidence interval 1.52 – 3.39, p = 4.9 e -5) from single institution retrospective cohort. **b)** as in (**a**) but depicting all lines of treatment; treatment line and endocrine therapy type (fulvestrant vs aromatase inhibitor) are included as a covariates in the multivariate model (hazard ratio 2.02 (1.52 – 2.70), 1.5 e -6). **c)** Progression free survival of patients treated with first-line CDK4/6i plus endocrine therapy combinations by g*BRCA2* status from the multi-institutional external validation cohort (hazard ratio 1.6 (1.3 – 2.0), p < 0.01). **d)** Swimmer’s plot depicting outcomes of patients receiving PARP inhibitors (green bar) after CDK4/6i and endocrine therapy (blue bar). Cox proportional hazard modeling of treatment type, stratified by patient ID approached statistical significance: HR 0.50 [0.24-1.03], p = 0.061. Measured interim response (defined as either CR; complete response or PR; partial response) on CDK4/6i vs. PARPi was assessed with Fisher’s exact test; p = 0.00078. OR 6.74, 95% CI: 1.93- 28.00). **e,f)** Two representative cases from the above plot are highlighted, denoting short progression free interval (7.3 mos and 3.3 mos, respectively) on first-line CDK4/6i and hormonal blockade. Both patients experienced a complete metabolic response on subsequent line PARP inhibitors and have either remained on this after 34.8 mos or progressed after 34.5 mos, respectively. **g)** Results of paired pre- and post-CDK4/6i sample analysis by germline and somatic *BRCA2* status, as well as germline *HRD*-related gene status. Oncoprint of post-CDK4/6i samples are depicted; cases where variant is not present in the pre-treatment sample outlined with black rectangle. Fisher exact test of *RB1* loss-of-function mutation acquisition by g*BRCA2* status: (OR 9.55, 95% CI 2.46 – 34.9, p = 0.00053.

We next validated our findings using an independent, nationwide clinicogenomic dataset containing manually curated patient-level outcomes data from both community oncology settings and academic medical centers (Flatiron Health database)^46^. This analysis confirmed a strong association between g*BRCA2* and shorter PFS on CDK4/6i plus ET (median PFS 12.0 vs 18.3 months, HR 1.83, 95% CI 1.36-2.48, p< 0.0001) (**Fig. 2c**) among the 2,185 patients with HR+/HER2- MBC who were treated with 1^st^ line CDK4/6i plus ET combinations.

We further investigated the MSK breast cancer cohort focusing on HR+/HER2- MBC patients who received a PARPi after progression on CDK4/6i (n = 35). We observed that HRD-directed therapy generally resulted in superior outcomes (**Fig. 2d**), with a trend towards improvement in PFS with PARPi compared to the previous CDK4/6i regimen (HR 0.50, 95% CI: 0.24-1.03, p = 0.061). Evaluating the patients who did not discontinue CDK4/6i or PARPi due to toxicity (n=34) allowed for more direct comparison of the clinical responses with either agent. PARPi, despite administration in a later line of therapy, produced a partial or complete response in 85.3% (n=29) of patients, whereas only 44.1% (n=15) of patients had a response to CDK4/6i and ET. Notably, the response to PARPi was more pronounced among the patients with no prior response to CDK4/6i and ET (Fisher’s exact test; p = 0.00078. OR 6.74, 95% CI: 1.93- 28.00).

Representative cases of rapid progression through first line CDK4/6i and ET, followed by prolonged complete metabolic response to PARPi are highlighted in **Fig. 2e,f**. Taken together, these results provide rationale for prioritization of PARPi for this genomically defined subgroup of breast cancer patients with expected poor outcomes on CDK4/6i-based combinations.

### Clinicogenomic analysis suggests evolutionary acquisition of *RB1* loss-of-function

To investigate the implications of germline status on the development of *RB1* loss, an established mechanism of CDK4/6i resistance, we assembled a large cohort of patients with paired tumor samples collected pre- and post-CDK4/6i therapy (n=541 tumors from 240 patients). Genomic analysis of this paired pre- and post-disease progression cohort demonstrated that acquisition of somatic *RB1* loss-of-function alteration was significantly more prevalent in *gBRCA2*-associated tumors vs g*BRCA2* WT tumors (31.6% vs 4.5%, OR 9.55, 95% CI 2.46 – 34.9, p = 0.00053) (**Fig. 2g**). The clinical responses and evolutionary trajectories of patients treated with CDK4/6i are highlighted by two representative g*BRCA2* HR+/HER2- metastatic breast cancer patients (**Fig. S9a,b**). Remarkably, acquired polyclonal *RB1* loss of function variants were seen in both cases in tumors collected post-CDK4/6i therapy. This convergent evolution under the selective pressure of CDK4/6i-based combinations resulting in the same mechanism of resistance in different tumor clones serves as evidence of strong and specific evolutionary pressure for *RB1* loss-of-function mutations as a dominant mechanism of CDK4/6i resistance in these g*BRCA2* tumors.

### Pretreatment copy number structure of *BRCA2*-driven tumors facilitates on-treatment *RB1* biallelic loss

Consideration of the genomic structure of *BRCA2-*driven tumors provides further insight into the development of *RB1* loss-of-function variants in *gBRCA2*-associated tumors. *BRCA2* and *RB1* are syntenic on chromosome 13q, and previous work suggests that biallelic inactivation of

*BRCA2* in g*BRCA2*-driven tumors often occurs through LOH resulting from loss of a large chromosomal segment inclusive of the WT allele^20,21^. To confirm that the pattern of *RB1* LOH was influenced by *gBRCA2* status in our cohort, we analyzed the patterns of *BRCA2* and *RB1* LOH, demonstrating a co-occurrence of *BRCA2* and *RB1* LOH in g*BRCA2* tumors compared to g*WT* (n = 63, 84.0% of evaluable cases; OR 11.5, 95% CI 6.08–23.63, p < 0.0001, **Fig. 3a**).

**Fig. 3:**
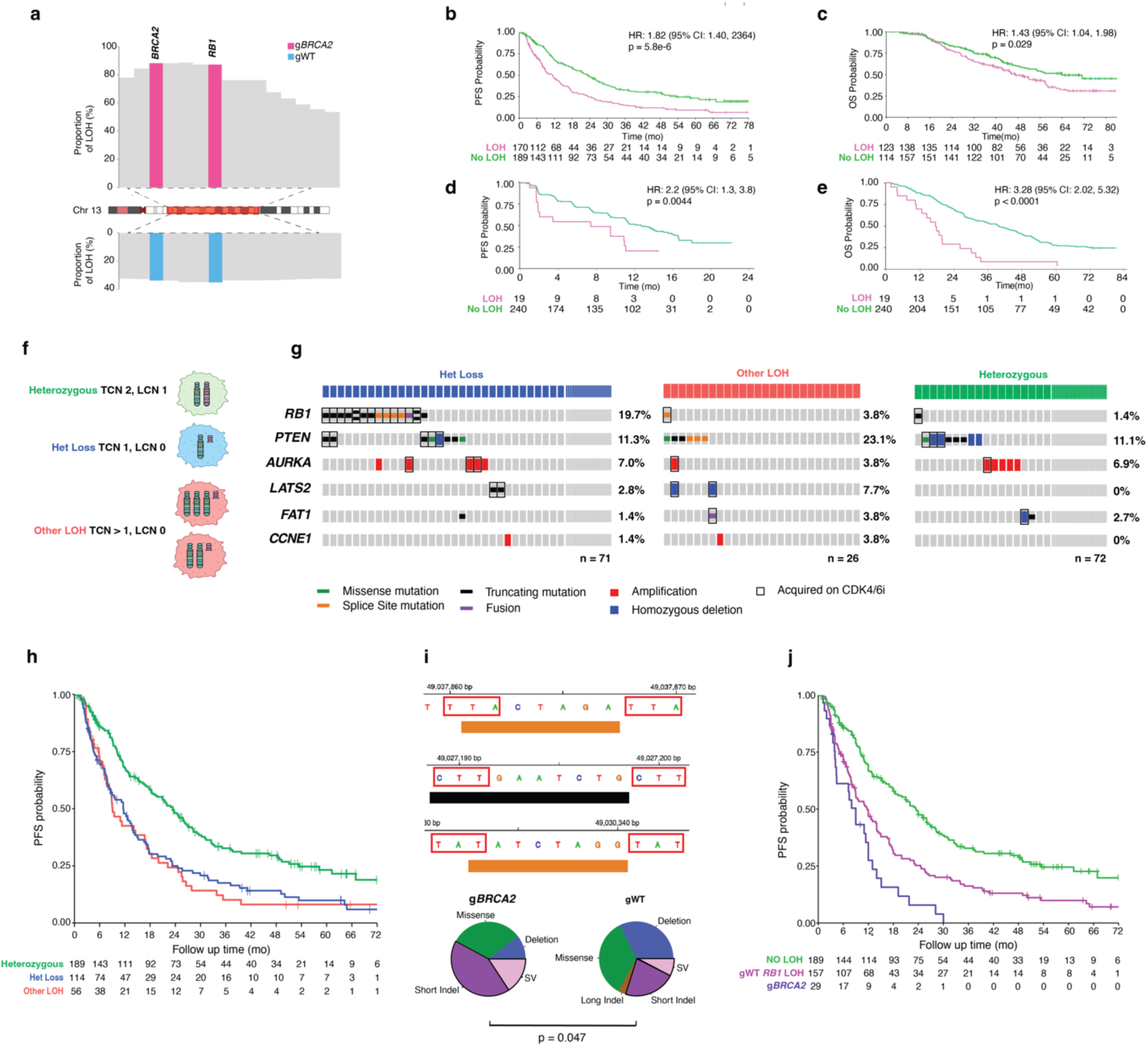
Dual effect of RB1 LOH and HRD mutagenesis on progression free survival and development of resistance mechanisms to CDK4/6i a) Comparison of LOH frequency of chromosome 13q in g*BRCA2* pathogenic variants (top), compared to g*WT* (bottom). The same segment spans *RB1* and *BRCA2* in 84.0% (n = 63) of evaluable g*BRCA2* cases, while a similar phenomenon is seen in wild-type cases in 31.3% (n = 527). Fisher test confirms that co-occuring *RB1* and *BRCA2* LOH is enriched in the g*BRCA2* cases (OR 11.5, 95% CI 6.08-23.63, p < 0.0001). All tumors included are from MSK breast cancer patients classified as HR+/HER2-. **B)** Progression free survival for HR+/HER2- MBC patients treated with first-line CDK4/6i plus ET by pre-treatment *RB1* LOH status, from the MSK Breast Cancer cohort. Multivariate Cox- proportional hazard ratio, including endocrine therapy partner as covariate (HR 1.82, 95% confidence interval 1.40 – 2.36, p = 5.8 e -6). **C)** as in **b**, but reporting overall survival (OS). The OS time is left-truncated to prevent immortality bias from the time of metastatic recurrence to the time of inclusion on the prospective sequencing protocol. Hazard ratio for OS 1.43 (1.04 – 1.98), p =0.029. **d)** PFS results from sequencing of baseline circulating tumor DNA from PALOMA-3 by baseline *RB1* LOH; HR 2.20 (95% CI 1.26 – 3.82), p = 0.0055. **e)** as in **d**, but depicting overall survival; HR 3.28 (95% CI 2.02 – 5.32), p < 0.001. **f)** Schematic representation of various total copy number (TCN) and lesser copy number (LCN) states that are included in the traditional definition of “*RB1* LOH*”*. This includes tumors not only with one total remaining allele (het loss, blue), but also cases with two or greater uniparental copies of the maternal or paternal allele (red). **g)** Results of paired pre- and post-CDK4/6i tumors analysis by pre-allelic status. Oncoprint of post-CDK4/6 samples are depicted. Cases where variant is not present in the pre-treatment sample outlined with black rectangle. Cases with more than one post-CDK4/6i variant in the same gene are indicated with two vertical variants. Fisher exact test of *RB1* acquisition by pre-treatment allelic configuration. Heterozygous *RB1* is the comparator: Het Loss: OR 15.7, 95% CI 2.23 – 683.3, p = 0.00050; Other LOH: OR 2.80, 95% 0.35 – 225.6, p = 0.46. **h**) Progression free survival of patients on first-line CDK4/6i plus ET combinations by refined allele-specific *RB1* copy number status, from the MSK Breast Cancer cohort. Multivariate Cox-proportional hazard ratio, including endocrine therapy partner and age at metastatic diagnosis as covariates; HR of RB1 het loss group: 1.76 (95% CI 1.32 – 2.34), p = 0.00011; HR of *RB1* other LOH = 1.95 (95% CI 1.37 – 2.77), p = 0.00020). **i)** Left: Nucleotide context of select *RB1* variants in g*BRCA2*-associated tumors, depicting short indel flanked by regions of microhomology. Right: Comparison of putative HRD-associated somatic variants of *RB1* (short indels, defined as < 25 bp, and structural variants) by g*BRCA2* status. Fisher exact test: p = 0.047 (OR 2.3, 95% CI 1.01 - 5.3).**j)** Progression free survival of patients treated with first-line CDK4/6i plus ET by *RB1* LOH status and *gBRCA2* status. Pre-treatment *RB1* LOH in gWT patients was associated with a significantly shorter PFS in gWT cases (Multivariate HR 1.84, 95% CI 1.44–2.37, p = 2.7e-6). Germline *BRCA2* status conferred an even shorter PFS (Multivariate HR 3.47, 95% CI 2.26 – 5.33, p = 1.3e-8). The above values were adjusted for endocrine therapy partner..

Despite this enrichment for *RB1* LOH in g*BRCA2* tumors, it was notable that *RB1* LOH was also observed in the larger gWT subset (n = 527/1768, 31.3% of evaluable cases). We next validated this finding using an external cohort by performing allele-specific copy number analysis on 46 g*BRCA2* cases which were profiled with whole exome sequencing (WES cohort, 24 patients from the University of Pennsylvania Abramson Cancer Center and 22 from Mayo Clinic).

Consistent with the MSK breast cancer cohort, 82.6% (n=38) of tumors in this WES cohort demonstrated concurrent LOH of *BRCA2* and *RB1* (p < 0.0001, **Fig. S10**).

Based on our *BRCA2:RB1* co-LOH results, we hypothesized that *RB1* LOH was the predominant mechanism of CDK4/6i resistance in g*BRCA2* associated tumors. Hence, we extended our survival analyses to consider whether pre-treatment *RB1* zygosity was predictive of clinical benefit from CDK4/6i therapy beyond the context of g*BRCA2* status. Specifically, we analyzed HR+/HER2- MBC patients in the MSK breast cancer cohort who underwent genomic profiling of tumors collected prior to start of 1^st^ line CDK4/6i therapy from which LOH status was evaluable (n=365 patients). *RB1* LOH was observed in 47.4% of these patients and was predictive of both significantly shorter PFS (HR 1.82, 95% CI 1.40 – 2.36, p = 5.8e-6), as well as shorter overall survival (OS, HR 1.43, 95% CI 1.04 – 1.98, p = 0.029, **Fig. 3b-c**).

To confirm these findings, we performed an analysis of pre-treatment cell-free DNA (cfDNA) samples collected as part of the PALOMA-3 study^22,23^, the pivotal randomized phase III trial of palbociclib plus fulvestrant versus fulvestrant monotherapy in patients with HR+/HER2- MBC. Baseline cfDNA samples from the palbociclib combination arm were sequenced using a 1729 amplicon custom AmpliSeq panel, which included 119 SNPs located within the *RB1* gene.

Circulating tumor DNA-based LOH analysis was conducted using a bespoke pipeline^23^, which identified *RB1* LOH in 19 (7.3%) of 259 pre-treatment cfDNA samples. *RB1* LOH was observed less frequently in the PALOMA-3 cfDNA cohort than in the MSK breast cancer cohort, likely due to the lower tumor fraction in cfDNA and lower sensitivity for detection of LOH in cfDNA compared to tumor tissue genomic data. Despite these technical limitations, *RB1* LOH in the PALOMA-3 cohort was associated with a significantly shorter PFS (HR 2.2, 95% CI 1.3 – 3.8, p = 0.0044), as well as an inferior OS (HR 3.28, 95% CI 2.02 – 5.32, p < 0.0001, **Fig. 3d,e**).

### Pre-treatment *RB1* allele-specific copy number profile predicts acquisition of *RB1* loss under the therapeutic pressure of CDK4/6i combination therapy

We hypothesized that the number of functional pre-treatment *RB1* alleles might influence the likelihood of acquiring an *RB1* loss-of-function mutation under the therapeutic pressure of CDK4/6i-based combination therapy. Specifically, we hypothesized that tumors with only one functional *RB1* copy would be statistically more likely to develop an *RB1* loss-of-function mutation as a second hit, leading to biallelic *RB1* loss conferring primary therapeutic resistance, as compared to those with two or more functional copies, which require multiple somatic events (hits) to achieve complete *RB1* loss. Hence, we investigated the clinical relevance of heterozygous *RB1* loss (a single allele of *RB1*), as compared to other allelic configurations such as loss before or after whole genome duplication resulting more than one remaining WT *RB1* allele (**Fig. 3f**) and extended our analysis of pre- and post-CDK4/6i tumors to 169 patients with evaluable data for these refined *RB1* allele-specific copy number definitions. Consistent with our hypothesis, only pre-treatment *RB1* het loss uniquely predisposed to acquisition of *RB1* loss-of-function upon exposure to CDK4/6i as compared to *RB1* diploid status or other LOH allelic configurations (p = 0.0061, **Fig. 3g**). Indeed, acquired *RB1* loss-of-function alterations were observed almost entirely in tumors with pre-treatment *RB1* heterozygous loss (87.5%), whereas such alterations were rarely observed in other settings.

To distinguish between predictive versus prognostic properties of pre-treatment *RB1* copy number changes, we repeated the outcome analyses based on the redefined pre-treatment *RB1* allelic state. Heterozygous loss was present in 31.8% (n=114) of pre-treatment tumors, whereas 14.6% (n=56) demonstrated other LOH configurations. Both heterozygous loss and other LOH allelic configurations were associated with worse outcomes on CDK4/6i-based combinations (**Fig. 3h**). As chromosomal instability is a known poor prognostic factor in both the pan-cancer and breast cancer settings^24^, we repeated this analysis adjusting for fraction genome altered (FGA) and whole genome doubling (WGD) as a covariate (**Fig. S11)**. Notably, *RB1* heterozygous loss was the sole significant allelic configuration associated with shorter PFS in this multivariate analysis, whereas other *RB1* LOH patterns were no longer associated with shorter PFS when adjusted for relevant genomic covariates (**Fig. S12a**). Conversely, an overall survival analysis with refined allele-specific copy number assessments demonstrated that allelic imbalances other than RB1 heterozygous loss were uniquely associated with a worse prognosis (HR 1.82, 95% CI 1.19 – 2.79, p = 0.0057, **Fig. S12b**). These results suggest that the grouping of other allelic imbalance patterns in *RB1* represents a *prognostic* biomarker, whereas *RB1* heterozygous loss serves as a *predictive* biomarker of outcome to CDK4/6i therapy specifically.

Although the acquisition of *RB1* loss-of-function variants during treatment with CDK4/6i is relatively rare in an unselected population^25,26^ (7.5% in our paired pre- and post-CDK4/6i treated cohort), our results demonstrate that acquired *RB1* loss is prevalent, and almost exclusively observed in tumors with only one functional allele of *RB1* prior to treatment initiation. This finding has major clinical implications for the development of novel surveillance strategies and for patient selection for clinical trials of precision therapeutic strategies directed against complete *RB1* loss^27,28^. These findings also hold potential clinical importance beyond *RB1*, as our results suggest that the pre-treatment allelic structure of tumor suppressor genes can predict not only the therapeutic outcome of specific targeted therapies but could also be used to predict the most likely *specific* mechanism by which the therapy will ultimately fail.

### HRD-driven mutagenesis facilitates acquisition of specific resistance mechanisms

We next assessed whether *gBRCA2*-mediated HRD specifically contributes to acquisition of somatic *RB1* loss-of-function variants during CDK4/6i therapy. Hence, we analyzed the types of alterations in *RB1* found in our g*BRCA2* cohort. We observed a statistically significant enrichment for short indels and structural variants within the coding sequencing of *RB1* in the g*BRCA2* setting, indicative of *BRCA2*-deficiency mediated lack of competent DNA repair (**Fig. 3i**). Indeed, the majority (75%, n=6) of post-CDK4/6i somatic *RB1* variants identified in g*BRCA2* tumors were short indels which predominantly exhibited the characteristic pattern of microhomology flanking a short indel, implicating a back-up DNA repair mechanism though microhomology-mediated end joining (MMEJ) in HRD-driven tumors^43,44^. Integration of our genomic analysis with treatment response data also suggested that HRD was likely responsible for the exceptionally rapid progression observed in g*BRCA2* carriers treated with CDK4/6i-based combinations. While *RB1* LOH was associated with a significantly shorter PFS in gWT cases (HR 1.84, 95% CI 1.44–2.37, p = 1.7e-6); patients with g*BRCA2* experienced a particularly poor PFS (HR 3.47, 95% CI 2.26 – 5.33, p = 1.3e-8) (**Fig. 3j**).

Thus, g*BRCA2* breast cancers with *RB1* LOH appeared to exploit the imperfections of back-up DNA repair mechanisms to escape therapeutic pressure. This phenomenon resembles the concept of “exaptation”, which refers to an evolutionary co-option of a trait to serve a purpose distinct from its original role^29^. Consequently, we expanded our analysis to assess the role of HRD (whether germline or somatic in origin) and clinical outcomes on CDK4/6i. We found that the presence of HRD-associated mutational signature in pre-treatment tumors of HR+/HER2- MBC patients conferred a significantly worse outcome on CDK4/6i plus ET combinations (HR 1.49, 95% CI 1.04 – 2.12, p = 0.030), even when analyzed outside the context of g*BRCA2* (**Fig S13**).

In sum, the data suggest that both an increased prevalence in *RB1* LOH and ongoing HRD-driven mutagenesis, which hastens the acquisition *RB1* loss-of-function mutations, contribute to the poor outcomes of breast cancer patients with *gBRCA2* mutations and HRD-driven tumors more broadly treated with CDK4/6i plus ET combinations.

## DISCUSSION

We describe disease and subtype-specific patterns of germline-somatic interactions and their clinical implications in a large clinico-genomic cohort of breast cancer patients. g*BRCA2* carriers experience a significantly worse outcome on combination CDK4/6i plus ET, the standard-of-care frontline approach in the metastatic HR+/HER2- setting. This finding was validated using a large, independent, multi-institutional real-world dataset that was representative of the general population of MBC patients treated in community and academic settings. We demonstrate a proclivity for *gBRCA2* tumors to develop *RB1* biallelic loss as a mechanism of resistance to CDK4/6i-based combination therapy and propose two separate characteristics of g*BRCA2* tumors that facilitate the acquisition of *RB1* loss. First, g*BRCA2*-driven breast cancers are more likely to harbor *RB1* loss of heterozygosity prior to the start of CDK4/6i therapy; we demonstrate that this allelic configuration independently predisposes to loss of the unaltered *RB1* allele. Second, the mutagenic processes active in HRD-driven tumors promote the selection of *RB1* loss-of-function mutations as a mechanism of biallelic inactivation (a “second hit”).

These observations raise provocative questions regarding the optimal therapy sequence for g*BRCA2* driven breast cancers. PARPi represent a proven synthetic lethality-based treatment strategy for patients with HRD tumors^16,30,31^. However, given the long-standing use of endocrine therapy combinations as front-line standard-of-care for HR+ breast cancers, use of PARPi is typically relegated to a later line of therapy following the emergence of resistance to CDK4/6i plus ET. Despite this typical sequencing of therapies, our data reveal a potentially greater benefit for later-line PARPi over earlier CDK4/6i plus ET in this molecularly defined subset of breast cancer patients. Intriguingly, the dominant mechanism of resistance to CDK4/6i plus ET may reduce the utility of several therapies typically used later in the disease course that rely on an intact G1 checkpoint. As the development of *RB1* loss appears to be partially mediated by HRD, earlier use of a PARPi may suppress this specific route of drug resistance resulting in a longer duration of response to subsequent CDK4/6i combinations. Specifically, reversion mutations that restore homologous recombination proficiency represent a widely prevalent and established mechanism of resistance to PARPi in g*BRCA2* carriers^14,32^. Therefore, tumor cells with somatic reversion mutations in *BRCA2* may be less likely to develop HRD-driven, *RB1* loss-of-function mutations under the selective pressure of subsequent CDK4/6i therapy.

The insights gleaned from our analysis should prompt a re-evaluation of conventional clinical practice in which the sequence of therapies is dictated by the clinical benefit observed in a biomarker unselected population. Most relevant to our current study, our results raise the provocative question of whether treatment with a PARPi as first-line therapy in patients with pathogenic g*BRCA2* variants or HRD tumors more broadly would lead to overall better outcomes not only in the first line setting, but also establish the genomic prerequisites for more successful subsequent later lines of therapy. This hypothesis serves as the impetus for the EvoPAR Breast01 study (NCT assignment pending), a randomized international phase 3 clinical trial which will compare frontline Saruparib (a PARP1 selective inhibitor) plus camisestrant (an oral selective estrogen receptor degrader) versus standard-of-care CDK4/6i combination therapy in patients with HR+/HER2- advanced breast cancer harboring alterations in selected HRD genes.

More generally, we demonstrate that pre-treatment heterozygous loss of *RB1* predicts the development of a somatic loss-of-function variant involving the wildtype *RB1* allele under the selective pressure of CDK4/6i (**Fig. 4a**). This proposed mechanism mirrors Knudson’s two-hit model^7,9^, which seminally demonstrated that copy number loss occurs as a necessary “second hit” for oncogenesis in patients with hereditary retinoblastoma. In contrast, in our proposed model, preexisting *RB1* heterozygous loss serves as the “first hit” increasing the likelihood of developing biallelic *RB1* loss and CDK4/6i resistance via acquisition of *RB1* loss variants as the “second hit”. Further studies will be needed to better refine the genomic and transcriptomic contexts predictive of CDK4/6i-resistance resulting from biallelic *RB1* inactivation. Preclinical models suggest that monoallelic *RB1* loss states may be sufficient for telomere instability and epithelial to mesenchymal transition signatures^33,34^. However, multiple studies demonstrate that *RB1* LOH does not always correlate with reduced *RB1* protein expression^35–37^. Furthermore, if *RB1* het loss was universally sufficient for intrinsic CDK4/6i resistance, acquisition of post- treatment *RB1* variants would be evenly distributed amongst all queried pre-treatment allelic states.

**Fig. 4:**
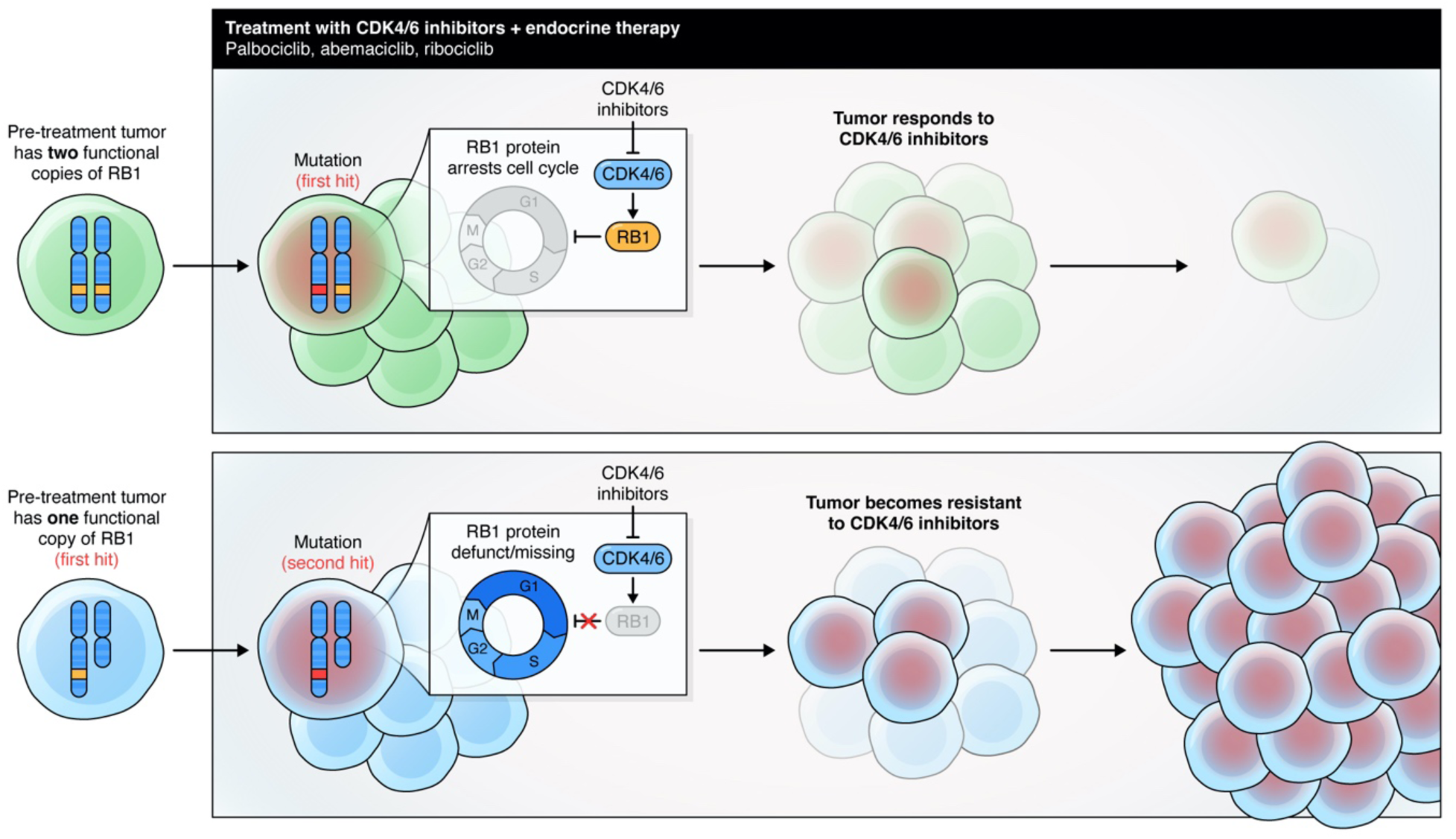
Proposed mechanism of “acquired second hit”. Our data demonstrates that pre-treatment allelic configuration of *RB1* influences treatment duration for breast cancer patients treated with CDK4/6i plus endocrine therapy combinations while also predicting for the precise mechanism of resistance.Tumors with only one allele present pre-treatment (*RB1* het loss or *RB1-*WT/*RB1-*loss, depicted in *blue)* experience complete *RB1* loss upon acquisition of a loss-of-function variant affecting the remaining *RB1* allele. Therefore, this clone is more likely to expand under the selective pressure of CDK4/6i and endocrine therapy, resulting in disease progression.

Our study has potential limitations. First, we focused our analysis specifically on exonic somatic variants that have previously demonstrated functional significance. This approach may fail to capture potentially consequential interactions, such as intronic variants of *RB1*, recently elucidated in a cohort of small cell lung cancer patients^38^. Second, given the real-world nature of this retrospective analysis, time between a particular event (i.e. start of treatment or disease progression) and a particular biopsy varied. Additionally, our paired pre- and post-CDK4/6i- treatment analysis may have failed to capture clonal expansion events that were extinguished by further therapy. Despite these caveats, we were able to conclusively demonstrate that pre- treatment *RB1* allelic configuration predicts for the development of a “second hit” resulting in loss-of-heterozygosity.

From a translational perspective, this work contributes to ongoing efforts to hasten the identification and interception of acquired resistance mechanisms. The ability to predict the precise mechanism of acquired resistance from genomic characteristics detectable in the pre-treatment sample would be a valuable tool in the precision oncology arsenal. Currently defined “targetable” genomic lesions encompass mutations, copy number homozygous deletions and amplifications, and structural variants which in their present genomic configuration have already disrupted a specific oncogenic pathway. With this work, we expand the paradigm of the “actionable genome” to states of allelic imbalance that have no biologic activity in their current allelic configuration but are predictive of therapeutic failure and thus could be used to guide early interception strategies designed to delay the onset of therapeutic resistance.

## METHODS

### Study cohort and prospective sequencing

The study cohort comprised 5,601 tumor samples from 4,640 breast cancer patients. All patients underwent prospective clinical tumor and germline sequencing as part of their clinical care (February 2014 to July 2017). 1,886 of these patients were included in previous clinicogenomic cohorts^39^. The present study was approved by the Memorial Sloan Kettering Cancer Center (MSK) Institutional Review Board (IRB) and all patients provided written informed consent for tumor and paired germline sequencing and review of medical records for demographic, clinical and pathology information. Genomic sequencing was performed on tumor DNA extracted from formalin-fixed, paraffin-embedded tissue and normal DNA extracted from mononuclear cells from peripheral blood in all patients. Patient samples were sequenced in a CLIA-compliant laboratory using one of three versions of the MSK-IMPACT-targeted sequencing panel, which included 341, 410-gene or 468 genes depending on the assay versions with somatic analysis described previously^40,41^. Tumors were obtained from the primary site in 50.8% of patients and from a metastatic site in 49.2% of patients.

Germline variant calling was performed using a sequence analysis pipeline validated for clinical use in a CLIA-compliant laboratory performing clinical sequencing of patient tumors and matched normal blood specimens as a part of routine clinical care^42^. To identify pathogenic and likely pathogenic germline variants associated with increased cancer risk, we employed a machine learning-based methodology previously described in Srinivasan et al^11^. Briefly, this framework incorporated mutation type, functional gene class, population allele frequencies, and orthogonal in silico tools to predict functional impact, and was trained on a cohort of 6,009 patients with cancer whose germline data were prospectively curated by expert clinical molecular geneticists using American College of Medical Genetics guidelines for clinical interpretation.

This framework was applied to a cohort of breast cancer patients at our institution who underwent prospective clinical genomic profiling under our sequencing protocol from April 2015 to August 2019 (n = 4639). Patient demographic, pathologic (including subsequent tumor sequencing data) and detailed clinical information was collected until the date of data freeze (June 2022). Low-risk variants in CHEK2 such as p.Ile157Thr were excluded from the analysis.

A total of 1927 patients had consented for identified analysis of germline variants via an IRB-approved protocol (NT01775072). Genomic and limited clinical data for all other patients analyzed in the present study (n = 2695) were anonymized before germline variant discovery and subsequent analyses. For numerical variables (such as time on treatment), this anonymization entailed rounding to the nearest digit; treatment times were therefore rounded to the nearest month. All single-nucleotide variants and small insertions and deletions (indels) identified in any blood normal sample were annotated with myvariant.info (as of August 2017)^43^. Population frequencies were obtained from gnomAD (r2.0.2)^44^. Curated assessments for pathogenicity were obtained from ClinVar (as of September 2017)^45^.

### Zygosity inference

We inferred somatic zygosity for all germline pathogenic variants using locus- and allele-specific DNA copy number inference^46^. All allele specific copy number (ASCN) solutions (FACETS outputs) from tumors with germline pathogenic variants were manually reviewed to ensure that the optimal solution was selected. Purity estimates were similarly inferred from the FACETS. We incorporated ASCN, purity and variant allele frequency into a previously described (Jonsson et al) framework, allowing statistical inference of heterozygous, biallelic (loss of wild type allele), or loss of mutant allele. The zygosity of the germline variant was considered indeterminate and excluded from zygosity analyses if the: (1) variant was homozygous in the germline; (2) read depth of coverage in the normal blood specimen was <50; (3) FACETS-derived total and minor copy number were not evaluable at the corresponding locus, or (4) An optimal FACETS solution could not able to be identified, most commonly from due to low tumor purity. Using these criteria, out of the 386 total cases with germline PV in the genes of interest, somatic zygosity status was evaluable for 312 (80.8%) of tumors. We excluded cases where copy number analysis and variant allele frequency was suggestive of loss of mutant allele, rather than wild type (n = 27, 7.90%) from further analyses..

To initially determine whether a given germline variant was in allelic imbalance in the corresponding tumor specimen, we evaluated consistency between observed somatic VAF and expected VAF. The latter value was calculated as a function of allele-specific DNA copy number and purity as follows:

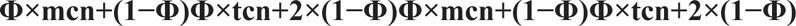

where *ɸ* is the tumor purity and tcn and lcn are the total and lesser copy number at the locus spanning the variant. Germline variants were considered heterozygous if their observed VAF was either (1) consistent with the expected VAF (within its 95% binomial CI) given balanced heterozygosity (tcn and mcn of either 2 and 1 or 4 and 2 in diploid and genome doubled tumors, respectively), or (2) less than the lower bound of the 95% CI of the expected VAF corresponding to a tcn and mcn of 3 and 1, respectively, which was either single copy gain of the mutant or WT allele. Germline variants in allelic imbalance of any kind were those with an observed VAF that was either within or greater than the 95% CI of the expected VAF corresponding to a copy-number state other than balanced heterozygosity^56^. For allelically imbalanced germline variants, loss of the WT was determined as those with an observed VAF within the 95% CI (or greater than the lower bound of the 95% CI) of the expected VAF corresponding to a lcn equal to 0 (observed VAF is concordant with the expected VAF when the lesser allele has a copy number of 0).

### Clinical data annotation

Detailed treatment history data were obtained for each of patients and included all lines of systemic therapy from the time of diagnosis to the study data lock (June 2022). The exact regimen, dates of start and stop therapy, as well as date of progression was annotated via expert chart review.

Each patient in the cohort was assigned a singular receptor status. Recognizing potential intertumoral heterogeneity, we sought a unified definition as follows: i) in cases where any metastatic biopsy was sequenced, receptor status was defined by treating clinician interpretation at the time of assigned first-line treatment; ii) in cases where only a primary tumor is sequenced, receptor status was defined by receptor status of the <u>sequenced</u> primary. We excluded cases where sequencing was obtained for external consultation (and therefore, lacked clinical or pathological data, n = 176), DCIS (n = 1), or cases where multi-site sequencing demonstrated no intra-patient genomic overlap and multiple distinct receptor statuses consistent with multiple primary tumors (n = 4) (**Fig. S1A)**. Employing these definitions, the MSK Breast Cancer cohort consisted of 3046 patients (68.3%) with hormone receptor positive, HER2-neu non-overexpressed (HR+/HER2-) receptor status, 781 patients (17.5%) with triple negative breast cancer (TNBC), with the remainder (638, 14.2%) HER2 amplified (HER2+).

Progression events were defined as i) a radiographic or clinical event prompting change in systemic therapy or recommendation for locally targeted radiation therapy; or ii) documented clinician impression detailing progression, after which there was documented patient or MD preference to continue same therapy. For samples who were not consented to our germline sequencing protocol (IMPACT part C), clinical outcomes were anonymized to avoid identification per institutional protocol. Outcome data (progression free survival, overall survival) was rounded to the nearest month. For samples who were consented to our institutional protocol, outcomes were not rounded.

### Enrichment Analysis

We compiled mutations, fusions, copy number alterations predicted to be functionally significant by the OncoKB precision oncology variant database^47^. All putative *RB1* homozygous deletions were manually re-reviewed prior to anonymization. Cases where the putative homozygous deletion spanned beyond the size of an amplicon (i.e. an event which would be interpreted as incompatible with tumor survival) were flagged.

In cases where a patient had multiple samples sequenced, we compiled the total somatic variants called from either the sequenced primary sample or all sequenced metastatic samples (omitting the primary), to avoid duplicate samples and to ensure that each set of variants was assigned either “primary” or “metastasis” as a covariate. Upon excluding samples with indeterminable receptor status (as described in the “clinical data annotation” section), 4459 patients were eligible for Firth-penalized logistic regression.

Receptor status was also defined on a “per-patient” as described above. Genes with alterations in less than 2% of patients were omitted from the analysis. For each remaining combination of germline gene X and somatic gene Y, we performed a Firth penalized logistic regression to account for the sparsity of the dataset. Receptor status and sample type (metastatic vs. primary) were included as covariates. The analysis was also repeated for each receptor status subtype, as well as repeated for biallelic vs mono-allelic germline variants. The latter step was only performed for samples where zygosity was evaluable (n = 4398). Putative p-values were adjusted for multiple hypothesis testing using the Benjamini-Hochberg method; values with q-value < 0.10 were deemed to be statistically significant.

### Allele-Specific Copy Number Definitions

In the MSK Breast Cancer cohort, somatic loss of heterozygosity events of *RB1* were defined based on manual FACETS review of pre-CDK4/6i treatment samples. Of 470 patients with pre first line CDK4/6i biopsies, 104 were removed due to: low purity or other technical limitations (such as “waterfalling” artifact), or paucity of heterozygous SNPs allowing for confident lesser copy number inference. *RB1* loss of heterozygosity was coarsely defined as lesser copy number of zero, irrespective of total copy number, while heterozygous state was any lesser copy number greater than or equal to one. In the PALOMA-3 cohort, RB1 LOH was defined as previously described in O’Leary et al Cancer Discovery.

In the MSK Breast Cancer cohort, we further separated the *RB1* LOH group into i) heterozygous loss (“Het Loss”, defined as a state with total copy number of one and lesser copy number of zero) and ii) other LOH (defined as LOH, but with total copy number greater than one). Fraction genome altered (FGA) was also calculated for each pre-treatment sample and defined as the fraction of log2 copy number variation (gain or loss) > 0.2 divided by the total size of the copy number profiled region.

### Mutational Signature Analysis

For MSK-IMPACT samples with at least 5 somatic SNVs, mutational signatures were computed using Signature Multivariate Analysis (SigMA), a tool extensively bench- marked for the analysis of formalin-fixed paraffin-embedded samples subjected to multigene panel sequencing, as previously described^48^. A dominant signature for each sample was determined based on the proposed category assigned by SigMA.

### Clinical Outcome Analysis

We determined the association between genomic alterations and PFS with disease progression on therapy with CDK4/6 inhibitors or patient death. Disease progression was defined as the date of the radiology study that prompted a change in systemic treatment, intervention with locally directed therapy (e.g. radiation therapy), or otherwise an annotation in the chart documenting progression of disease. We categorized CDK4/6 inhibitor regimens based on their endocrine therapy partner (aromatase inhibitor vs selective estrogen receptor degrader/ SERD). Patients with ablation of known sites of disease with radiotherapy or surgical resection prior to initiation of CDK4/6i therapy were excluded, as were patients who discontinued therapy due to toxicity after two weeks (n=124). To maximize statistical power for downstream clinicogenomic analyses, we expanded the cohort to include CDK4/6i outcomes on all additional germline *BRCA2* pathogenic variants identified through our institutional germline pathogenic variant database (end date: 8/2022, resulting in 23 additional samples).

We employed both univariate and multivariate Cox proportional hazard models (stratified by endocrine therapy partner, and treatment line, where available). For patients with multiple lines of therapy from the same class of treatment, only the first treatment line from that class that was started after the MSK-IMPACT biopsy was included in the analysis. For analyses pertaining to allele-specific copy number, fraction genome altered and whole genome duplication were employed as additional covariates. These are recognized poor prognostic factors and may be a confounding factor given increased likelihood for tumors with measures of copy number instability to harbor LOH of any specific region.

For overall survival analysis, we implemented a left-truncated model to account of time from diagnosis of metastatic disease to enrollment on sequencing protocol. Similarly to the univariate analyses, we employed univariate and multivariate Cox proportional hazard models. In addition to endocrine therapy partner, age at metastatic diagnosis was also included as a covariate. We rejected the null hypotheses with a two-sided α = 0.05.

To accomplish the matched pairs analysis, we considered any paired pre-CDK4/6i and post- CDK4/6i treatment solid tumor samples sequenced on MSK-IMPACT. Allele-specific copy number analysis was performed for the pre-treatment cases; cases where low purity precluded confident allele specific copy number ascertainment were excluded. The exclusion of such low purity samples assures acquisition of post-treatment variants is due to tumor evolution rather than technical inability to capture the preceding variant.

For other exploratory analyses (comparison of sample type and receptor status by *RB1:BRCA2* co-mutation status, comparison of rates of RB1 loss-of-function sample acquisition by group, comparison of *RB1* short indels and SVs by *BRCA2* status, etc), details of statistical analysis can be found in the respective Fig. legend. Unless stated otherwise, statistical analysis of differences between samples was performed using two-tailed Fisher exact tests; p < 0.05 was defined as significant.

## DATA AVAILABILITY

The assembled prospective somatic and germline mutational data from tumors needed to repliacate our figures for the entire cohort are provided as Supplementary Tables. Deidentified clinical outcomes have been provided in the Supplementary Tables. Germline variants, tumor-specific zygosity, as somatic variants and allele specific copy number estimates for *RB1* will be made available from the National Center for Biotechnology Information dbGaP archive prior to publication. Otherwise, our analyses may be replicated with the following files provided:

**Supplemental Table 1 -** Anonymized germline pathogenic variant calls and corresponding anonymized somatic mutation, structural variant, copy number calls

**Supplemental Table 2 –** Anonymized demographics of dataset used for germline-somatic analysis

**Supplemental Tables 3-7** – Results of germline-somatic interactions for *BRCA2, BRCA1, ATM, CHEK2, PALB2*, respectively. This includes results of overall multivariate analysis, as well as grouped by receptor status and zygosity.

**Supplemental Table 8** – Anonymized outcomes of CDK4/6i and endocrine therapy. Additionally, curated description of progression free survival on PARPi subsequent to CDK4/6i in germline BRCA2 patients. Anonymized dataset denoting whether sample was collected before or after CDK4/6i. Lastly, summary of comparison of *RB1* variants in g*WT* compared to *g*BRCA2 setting.

**Supplemental Table 9** – Progression free survival and overall survival data corresponding to pre-treatment allele specific copy number analysis, including binary (LOH or no LOH) and refined allele-specific copy number assignment (Heterozygous, Het Loss, Other LOH) as determined by FACETS. Additionally, the data required to make the pre and post CDK4/6i Oncoprint including several implicated mechanisms of resistance (*RB1*, *PTEN*, *AURKA, LATS2, FAT1, YAP1, CCNE1)* is additionally provided.

## CODE AVAILABILITY

All code and scripts will be made available in a GitHub repository, which will describe how to reproduce the all figures and tables in this publication.

## DECLARATIONS

COI: P.R. has received institutional grant/funding from Grail, Novartis, AstraZeneca, EpicSciences, Invitae/ArcherDx, Biothernostics, Tempus, Neogenomics, Biovica, Guardant, Personalis, Myriad, Shares/Ownership interests in Odyssey Biosciences, and consultation/Ad board/Honoraria from Novartis, AstraZeneca, Pfizer, Lilly/Loxo, Prelude Therapeutics, Epic Sciences,Daiichi-Sankyo, Foundation Medicine, Inivata, Natera, Tempus, SAGA Diagnostics, Paige.ai, Guardant, Myriad. S.C. has received institutional grant/funding from Daiichi-Sankyo, AstraZeneca, and Lilly, Shares/Ownership interests in Odyssey Biosciences, Effector Therapeutics, and Totus Medicines, and consultation/Ad board/Honoraria from AstraZeneca, Lilly, Novartis, Neogenomics, Nuvalent, Blueprint, SAGA Diagnostics, and Effector Therapeutics. J.R.F, M.S, E.N, N.L., E.S. are currently employed by AstraZeneca. Y.L. and X.H. are currently employed by Pfizer. M.F.B. has received consultation from AstraZeneca, Eli Lilly and Paige AI, and research support from Boundless Bio, and declares intellectual property interests with SOPHiA Genetics.

## ACKNOWLEDGEMENTS

We gratefully acknowledge the following funding sources: Komen Career Catalyst Research Award, Gray Foundation (K.N., M.E.R.), BCRF (P.R, S.C., and K.N.), NCI P50 CA247749 (S.C.), ASCO Conquer Cancer Foundation YIA (A.S.), Basser Center for BRCA (K.N.)

## Supplemental Figure Legends

**Fig. S1) CONSORT diagram describing germline-somatic interaction analysis and CDK4/6i PFS analysis.** Consort diagram revealing case inclusion in germline somatic and CDK4/6i analysis comparing g*BRCA2* to g*WT.* Cases were excluded from further clinicogenomic analysis if no clinical or pathogical data was provided, if the patient did not have invasive breast cancer (i.e. ductal carcinoma in situ was sequenced), and if the patient had multiple evolutionarily distinct tumors with discordant receptor status (n = 181 excluded total). Additionally, patients were excluded from analysis involving zygosity for technical reasons (including indeterminate LOH in the genes of interest); patients with loss of mutated allele were also excluded from analysis. Patients were excluded from downstream analysis of CDK4/6i and endocrine therapy outcomes for the following reasons: i) if initial receptor status was interpreted to be TNBC or HER2+, ii) the patients received CDK4/6i for fewer than 2 weeks prior to toxicity, iii) if the known sites of disease were ablated prior to initiation of CDK4/6i and endocrine therapy (e.g., oligometastatic disease rendered to have “no evidence of disease” prior to initiation of systemic after radiotherapy).

**Fig. S2) Zygosity status of germline genes involved in homologous recombination pathway** Descriptive summary of zygosity status of each gene included in our study, including distinction between loss of wild type allele (“LOH_CN” or copy number loss, “Second Hit Mut.”) and loss of mutated allele (Loss Ref). Biallelic inactivation rates varied significantly across genes, ranging from 47.5% in g*CHEK2* carriers, to 88.8% and 89.7% for g*BRCA1* and g*BRCA2* carrier. As this is a descriptive figure, statistical analysis was not pursued.

**Fig. S3) Germline-somatic interactions of g*ATM*, stratified by zygosity and receptor status** Depiction of enrichment analysis of gATM compared to gWT. Overall results are depicted from iteratively conducting a Firth penalized regression overall all genes in which somatic variant is greater than 3% in the cohort. Model receptor status and zygosity were employed as covariates; results denote p-value as well as 95% confidence interval of odds ratio. Overall analysis was followed by stratification by zygosity and receptor status. For each scenario, q value was calculated. Significant somatic-germline interactions were defined by q < 0.10.

**Fig S4) Germline-somatic interactions of g*BRCA1*, stratified by zygosity and receptor status.** Depiction of enrichment analysis of g*BRCA1* compared to gWT. Overall results are depicted from iteratively conducting a Firth penalized regression overall all genes in which somatic variant is greater than 3% in the cohort. Model receptor status and zygosity were employed as covariates; results denote p-value as well as 95% confidence interval of odds ratio. Overall analysis was followed by stratification by zygosity and receptor status. For each scenario, q value was calculated. Significant somatic-germline interactions were defined by q < 0.10.

**Fig S5) Germline-somatic interactions of g*BRCA2*, stratified by zygosity and receptor status.** Depiction of enrichment analysis of g*BRCA2* compared to gWT. Overall results are depicted from iteratively conducting a Firth penalized regression overall all genes in which somatic variant is greater than 3% in the cohort. Model receptor status and zygosity were employed as covariates; results denote p-value as well as 95% confidence interval of odds ratio. Overall analysis was followed by stratification by zygosity and receptor status. For each scenario, q value was calculated. Significant somatic-germline interactions were defined by q < 0.10.

**Fig S6) Germline-somatic interactions of g*CHEK2*, stratified by zygosity and receptor status.** Depiction of enrichment analysis of g*CHEK2* compared to gWT. Overall results are depicted from iteratively conducting a Firth penalized regression overall all genes in which somatic variant is greater than 3% in the cohort. Model receptor status and zygosity were employed as covariates; results denote p-value as well as 95% confidence interval of odds ratio. Overall analysis was followed by stratification by zygosity and receptor status. For each scenario, q value was calculated. Significant somatic-germline interactions were defined by q < 0.10.

**Fig S7) Germline-somatic interactions of gPALB2, stratified by zygosity and receptor status.** Depiction of enrichment analysis of g*PALB2* compared to gWT. Overall results are depicted from iteratively conducting a Firth penalized regression overall all genes in which somatic variant is greater than 3% in the cohort. Model receptor status and zygosity were employed as covariates; results denote p-value as well as 95% confidence interval of odds ratio. Overall analysis was followed by stratification by zygosity and receptor status. For each scenario, q value was calculated. Significant somatic-germline interactions were defined by q < 0.10.

**Fig. S8) Oncoprints of enriched somatic-germline interactions.** Oncoprint depicting the somatic alteration type on a per-patient (column) and per-gene (row) basis. Characteristics of mutations, copy number alterations, and fusions in the indicated genes are noted; genes which have been determined to be significant on prior germline-somatic interaction analysis are included. Receptor status, sample type, and zygosity are annotated above. **a)** depicts Oncoprint from *gBRCA1* vs. gWT**, b)** depicts g*CHEK2* vs g*WT*, **c)** depicts g*ATM* vs g*WT***, d)** depicts g*PALB22* vs g*WT*

**Fig. S9) Representative cases of short PFS and evidence of convergent *RB1* evolution in g*BRCA2* treated with g*BRCA2* inhibitors. a)** Depicts a patient vignette of locally advanced ER+/PR-/HER2- breast cancer for which patient did not seek treatment for 2 years. She presented with progressive disease in bone and underwent SBRT to T8-T12. She was on CDK4/6i and aromatase inhibitor for 6 months, at which point she experienced oligo-progression in chestwall. These agents were continued for 90 more days, after which progression of disease was noted in parenchyma (liver, pleura, soft musculoskeletal tissue). She was subsequently plaed on fulvestrant, and again experienced progression of disease after 6 months. Post-progression plasma cell free DNA revealed 4 different *RB1* variants. The patient was subsequently treated with olaparib, to which a partial response was noted. She remained on this for 560 days until eventual progression in chest wall. Post progression next generation sequencing of the site of progression revealed a fifth *RB1* loss of functional variant which had not previously been noted. **b)** Depicts a patient vignette for locally advanced lobular carcinoma (ER+/PR+/HER2-) for which adjuvant chemotherapy was pursued. The patient developed distant metastatic recurrence after 34 months, after which she was placed on tamoxifen and ovarian suppression for 7 months. After subsequent progression of disease (chest wall, lymph nodes, ovaries), she was treated with second line CDK4/6i and aromatase inbhitor. She developed oligo progressive disease in the liver. Biopsy revealed a p.L452Yfs*14 variant in *RB1* which was predicted to be loss of function. Subsequent plasma cell-free DNA sequencing revealed an additional splice site variant in *RB1*.

**Fig. S10) Validation of concurrent *RB1* and *BRCA2* LOH in external gBRCA2 WES cohort.** FACETS was performed on whole exome sequencing of breast cancer samples from gBRCA2 Abramson Cancer Center at University of Pennsylvania (n = 24), and Mayo Clinic (n = 22). Of these samples, 38 (82.6%) demonstrated concurrent LOH of *BRCA2* and *RB1*. Two-sided fisher t-test between *RB1* LOH and *BRCA2* LOH was statistically significant (OR = *Inf*, 95% CI 11.12 – *Inf*, p = 3.0 e -6).

**Fig. S11) CONSORT Diagram describing pre treatment allele-specific copy number analysis.** Allele-specific copy number analysis was performed on tumor samples collected prior to initiation of first-line CDK4/6i and endocrine therapy. Samples were excluded (n = 140) for technical reasons including low purity or paucity of heterozygous SNPs in the region of interest. The consort diagram describes the clinical outcomes analysis on the 359 samples remaining, as well as the matched pairs analysis on the 169 samples with corresponding post progression biopsies.

**Fig. S12) Pre-treatment allele-specific copy number analysis. a.** Adjusted Kaplan-Meier curve, accounting for covariates including endocrine therapy partner, age at metastatic diagnosis, fraction genome altered (FGA), whole genome duplication (WGD). After adjusting for relevant genomic and clinical covariates, the group harboring “Other LOH” configurations of RB1 ceases to be associated with PFS (HR 1.35, 95% CI 0.91 – 1.99, p = 0.13). RB1 het loss remains associated with shorter PFS after these adjustments: HR 1.76, 95% CI 1.30 – 2.39, p = 0.00024). **b.** Overall survival curve of patients received first line CDK4/6i and endocrine therapy, based on pre-treatment RB1 allelic configuration. Age at metastatic diagnosis is included as a covariate. “Other LOH” is uniquely associated with a shorter overall survival: HR 1.82, 95% CI 1.19 – 2.79, p = 0.0057. HR of Het Loss: HR 1.32, 95% CI 0.92 – 1.90, p = 0.13).

**Fig. S13) Implications of HRD signature on CDK4/6i PFS in BRCA2 WT breast cancers**. Signature Multivariate Analysis (SigMA)^48^ was used to derive dominant mutational signature in available cases. Out of 412 cases of pre-CDK4/6i samples (across all treatment signs), 18 g*BRCA2* cases were excluded, and 188 cases were excluded as they did not yield a dominant mutational signature. Compared to non-HRD signature, a dominant HRD signature was associated with decreased progression free survival: HR 1.49, 95% CI 1.04 – 2.12, p = 0.030).

