## Supplemental Figures for "Tumor suppressor heterozygosity and homologous recombination deficiency mediate resistance to front-line therapy in breast cancer"

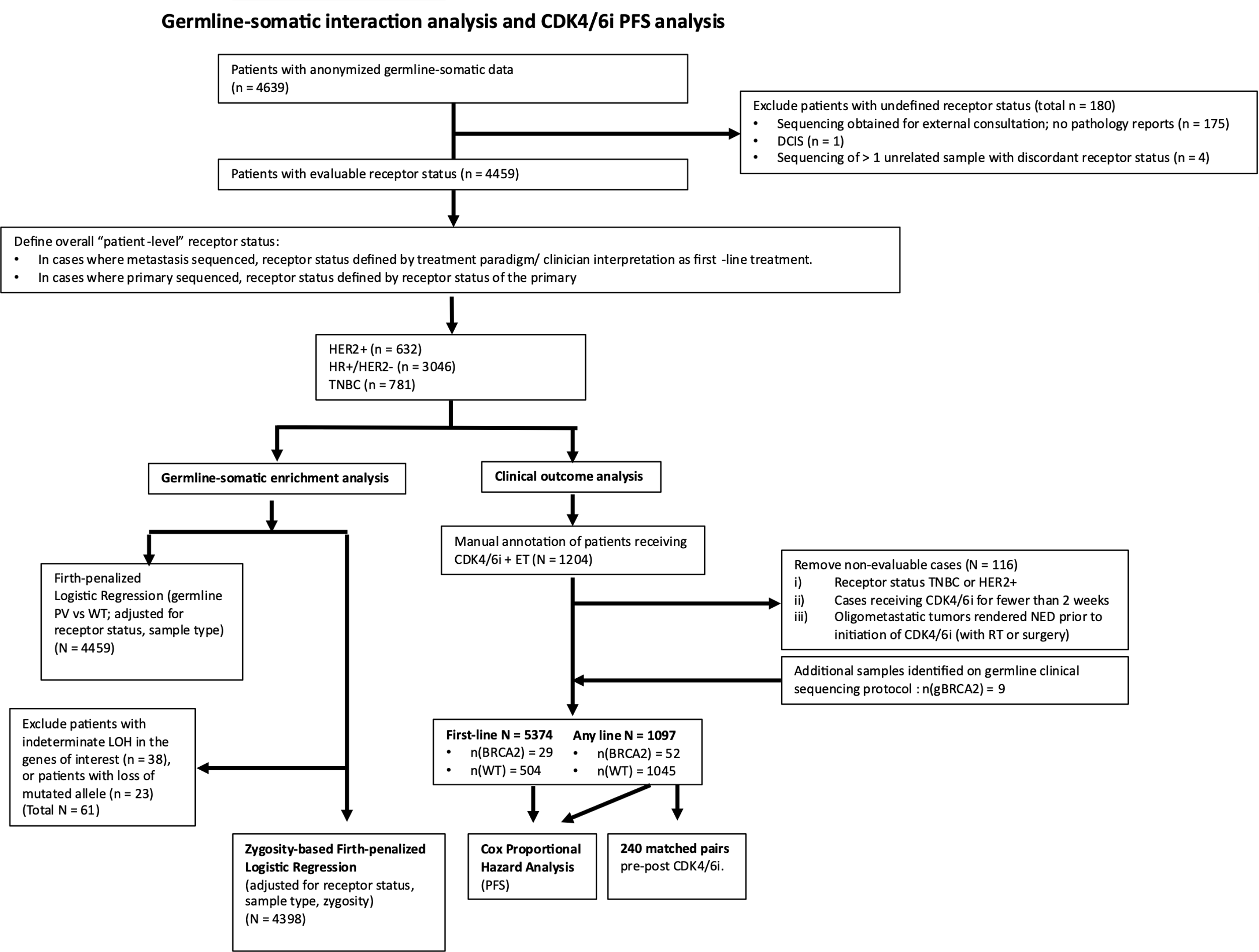

**Fig. S1) CONSORT diagram describing germline-somatic interaction analysis and CDK4/6i PFS analysis.** Consort diagram revealing case inclusion in germline somatic and CDK4/6i analysis comparing g*BRCA2* to g*WT.* Cases were excluded from further clinicogenomic analysis if no clinical or pathogical data was provided, if the patient did not have invasive breast cancer (i.e. ductal carcinoma in situ was sequenced), and if the patient had multiple evolutionarily distinct tumors with discordant receptor status (n = 181 excluded total). Additionally, patients were excluded from analysis involving zygosity for technical reasons (including indeterminate LOH in the genes of interest); patients with loss of mutated allele were also excluded from analysis. Patients were excluded from downstream analysis of CDK4/6i and endocrine therapy outcomes for the following reasons: i) if initial receptor status was interpreted to be TNBC or HER2+, ii) the patients received CDK4/6i for fewer than 2 weeks prior to toxicity, iii) if the known sites of disease were ablated prior to initiation of CDK4/6i and endocrine therapy (e.g., oligometastatic disease rendered to have “no evidence of disease” prior to initiation of systemic after radiotherapy).

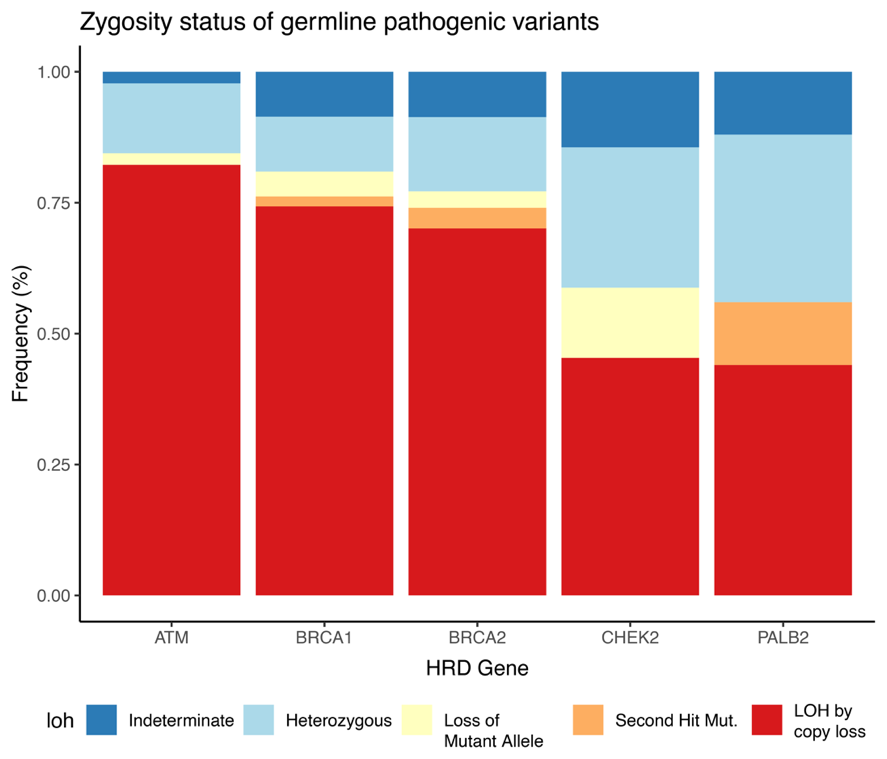

**Fig. S2) Zygosity status of germline genes involved in homologous recombination pathway**

Descriptive summary of zygosity status of each gene included in our study, including distinction between loss of wild type allele (“LOH_CN” or copy number loss, “Second Hit Mut.”) and loss of mutated allele (Loss Ref). Biallelic inactivation rates varied significantly across genes, ranging from 47.5% in g*CHEK2* carriers*,* to 88.8% and 89.7% for g*BRCA1* and g*BRCA2* carrier*.* As this is a descriptive figure, statistical analysis was not pursued.

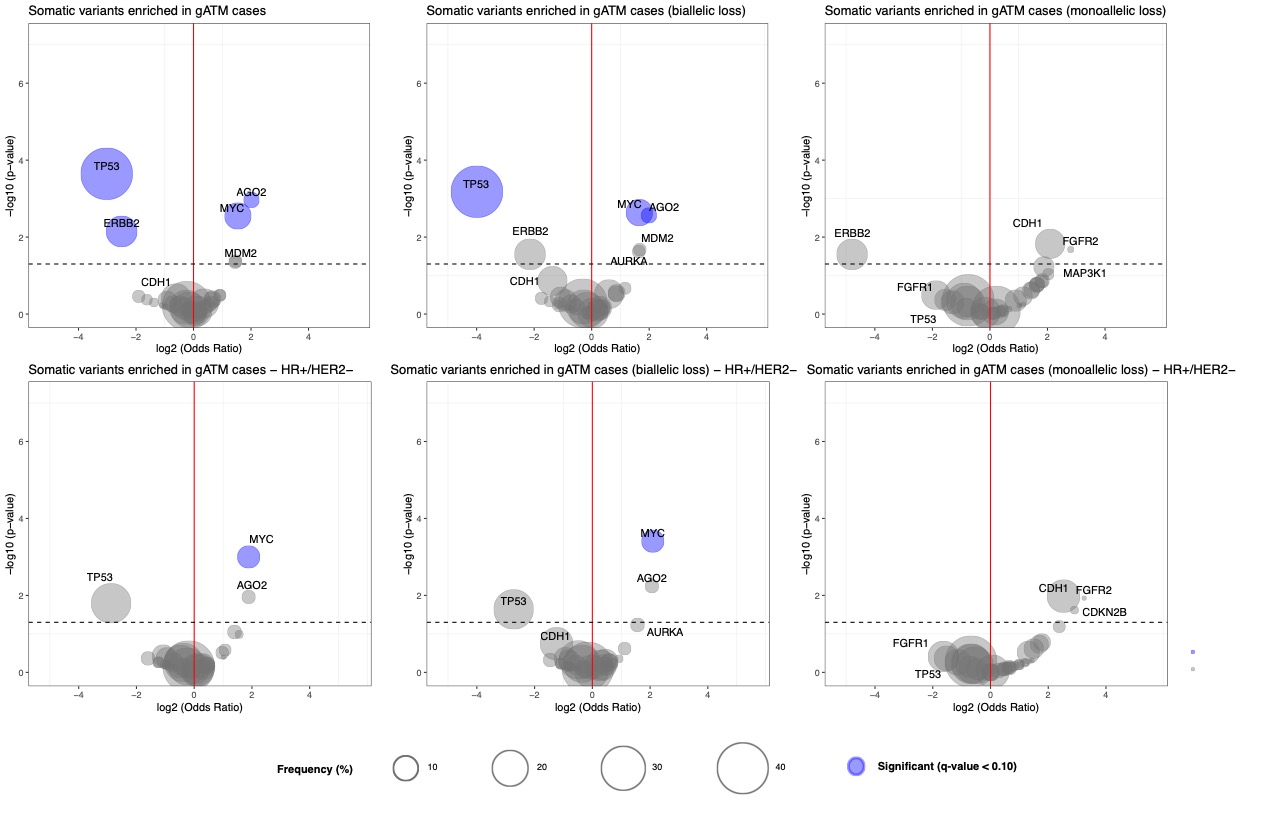

**Fig. S3) Germline-somatic interactions of g*ATM*, stratified by zygosity and receptor status**

Depiction of enrichment analysis of gATM compared to gWT. Overall results are depicted from iteratively conducting a Firth penalized regression overall all genes in which somatic variant is greater than 3% in the cohort. Model receptor status and zygosity were employed as covariates; results denote p-value as well as 95% confidence interval of odds ratio. Overall analysis was followed by stratification by zygosity and receptor status. For each scenario, q value was calculated. Significant somatic-germline interactions were defined by q < 0.10.

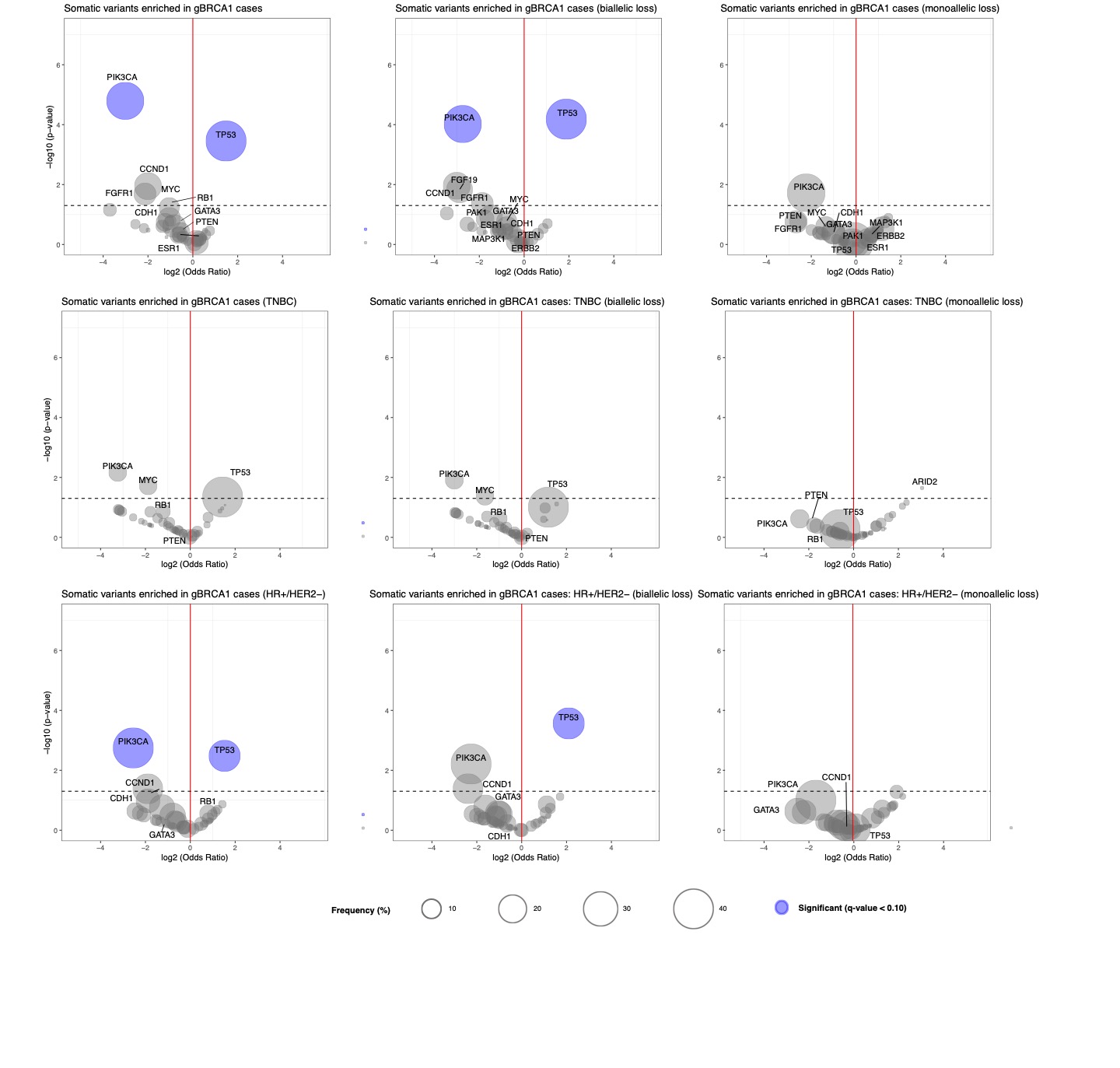
**Fig S4:** **Germline-somatic interactions of g*BRCA1*, stratified by zygosity and receptor status.** Depiction of enrichment analysis of g*BRCA1* compared to gWT. Overall results are depicted from iteratively conducting a Firth penalized regression overall all genes in which somatic variant is greater than 3% in the cohort. Model receptor status and zygosity were employed as covariates; results denote p-value as well as 95% confidence interval of odds ratio. Overall analysis was followed by stratification by zygosity and receptor status. For each scenario, q value was calculated. Significant somatic-germline interactions were defined by q < 0.10.

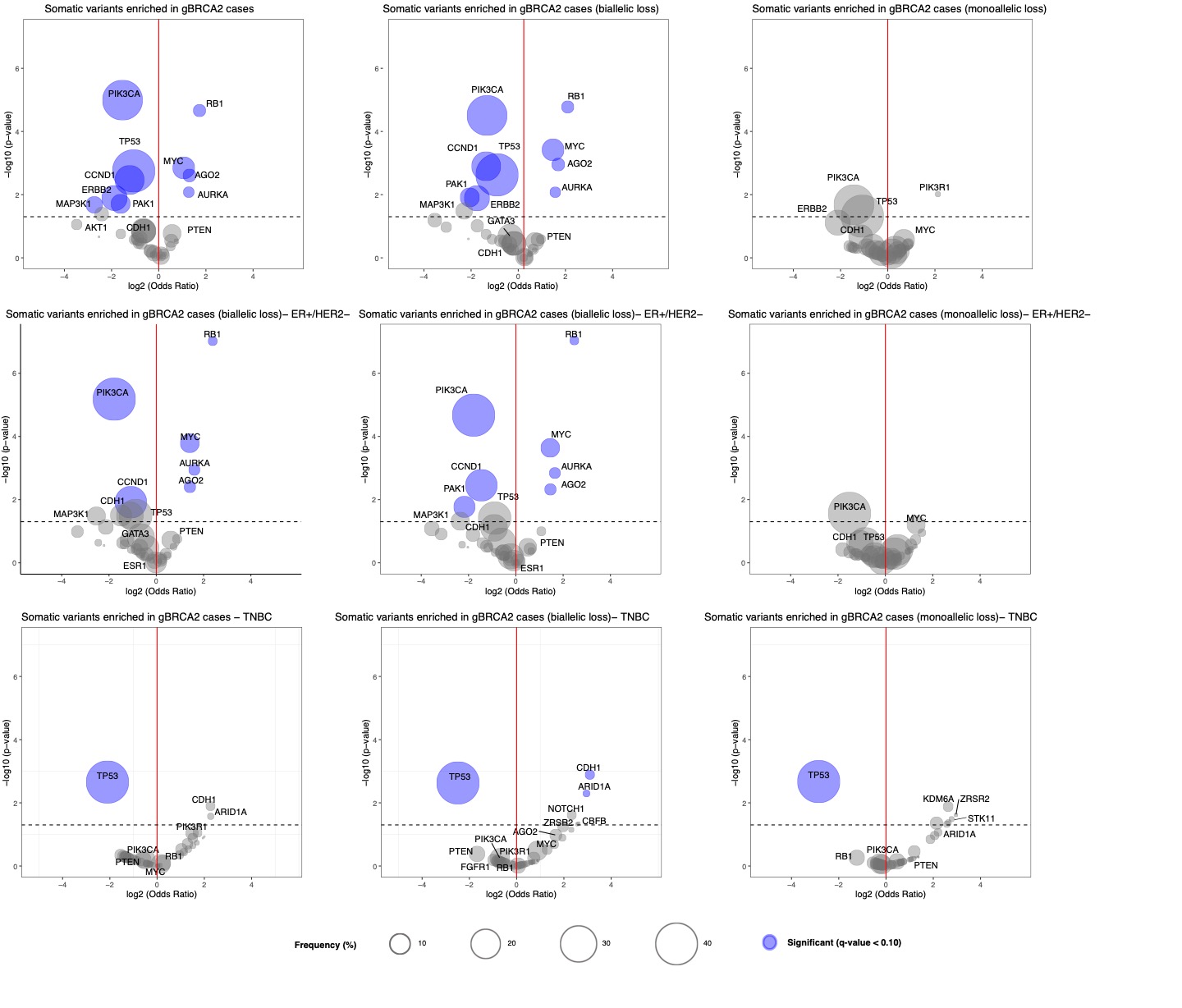

**Fig S5:** **Germline-somatic interactions of g*BRCA2*, stratified by zygosity and receptor status.** Depiction of enrichment analysis of g*BRCA2* compared to gWT. Overall results are depicted from iteratively conducting a Firth penalized regression overall all genes in which somatic variant is greater than 3% in the cohort. Model receptor status and zygosity were employed as covariates; results denote p-value as well as 95% confidence interval of odds ratio. Overall analysis was followed by stratification by zygosity and receptor status. For each scenario, q value was calculated. Significant somatic-germline interactions were defined by q < 0.10.

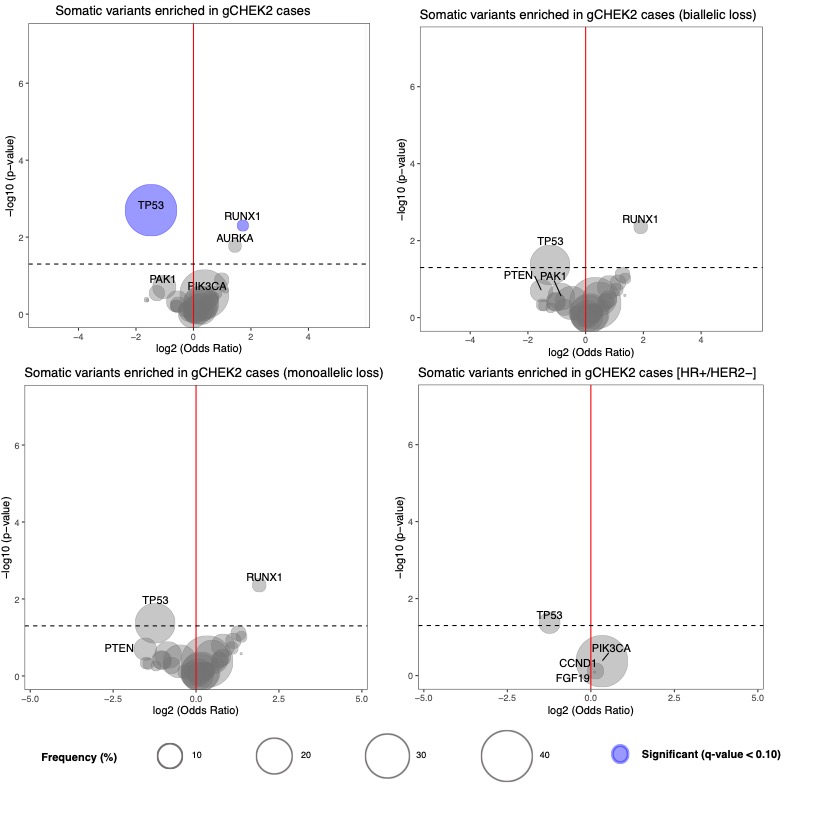

**Fig S6:** **Germline-somatic interactions of g*CHEK2*, stratified by zygosity and receptor status.** Depiction of enrichment analysis of g*CHEK2* compared to gWT. Overall results are depicted from iteratively conducting a Firth penalized regression overall all genes in which somatic variant is greater than 3% in the cohort. Model receptor status and zygosity were employed as covariates; results denote p-value as well as 95% confidence interval of odds ratio. Overall analysis was followed by stratification by zygosity and receptor status. For each scenario, q value was calculated. Significant somatic-germline interactions were defined by q < 0.10.

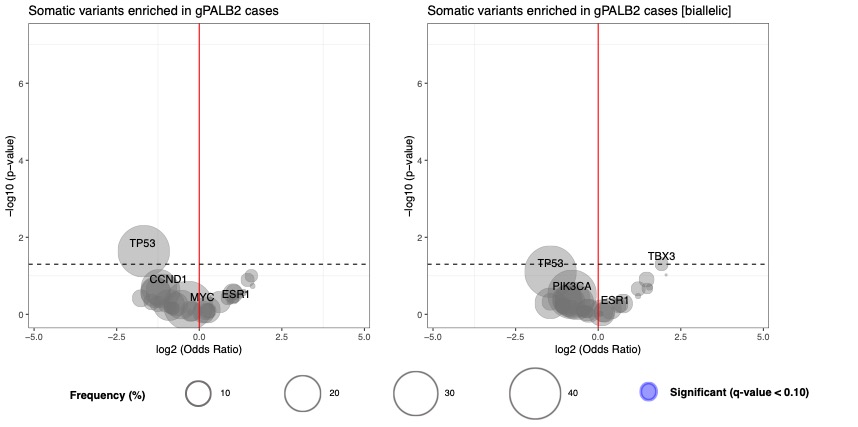

**Fig S7:** **Germline-somatic interactions of gPALB2, stratified by zygosity and receptor status.** Depiction of enrichment analysis of g*PALB2* compared to gWT. Overall results are depicted from iteratively conducting a Firth penalized regression overall all genes in which somatic variant is greater than 3% in the cohort. Model receptor status and zygosity were employed as covariates; results denote p-value as well as 95% confidence interval of odds ratio. Overall analysis was followed by stratification by zygosity and receptor status. For each scenario, q value was calculated. Significant somatic-germline interactions were defined by q < 0.10.

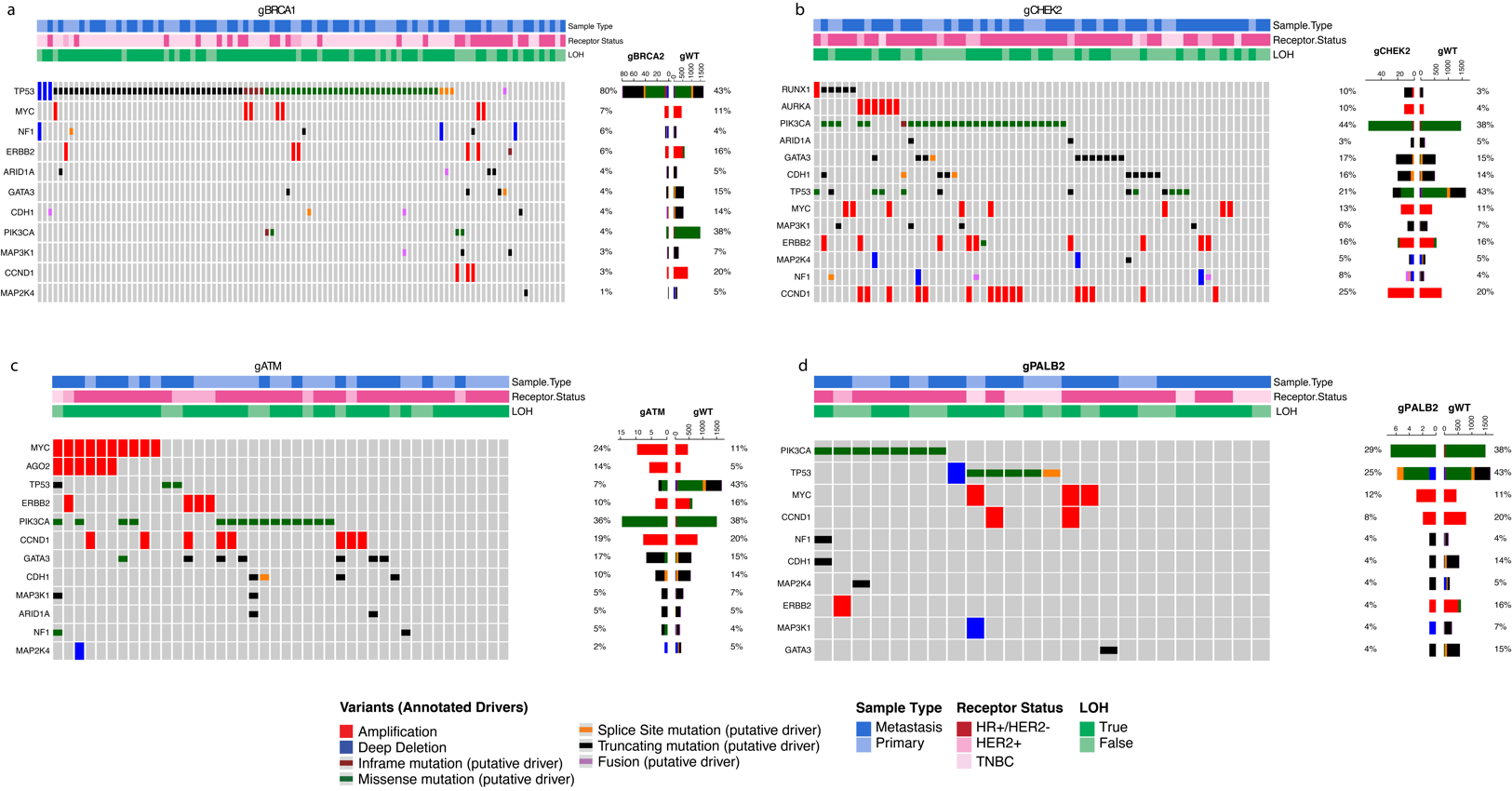

**Fig. S8) Oncoprints of enriched somatic-germline interactions.** Oncoprint depicting the somatic alteration type on a per-patient (column) and per-gene (row) basis. Characteristics of mutations, copy number alterations, and fusions in the indicated genes are noted; genes which have been determined to be significant on prior germline-somatic interaction analysis are included. Receptor status, sample type, and zygosity are annotated above. **a)** depicts Oncoprint from *gBRCA1* vs. gWT, **b)** depicts g*CHEK2* vs g*WT*, **c)** depicts g*ATM,* **d)** depicts g*PALB22* vs g*WT.*

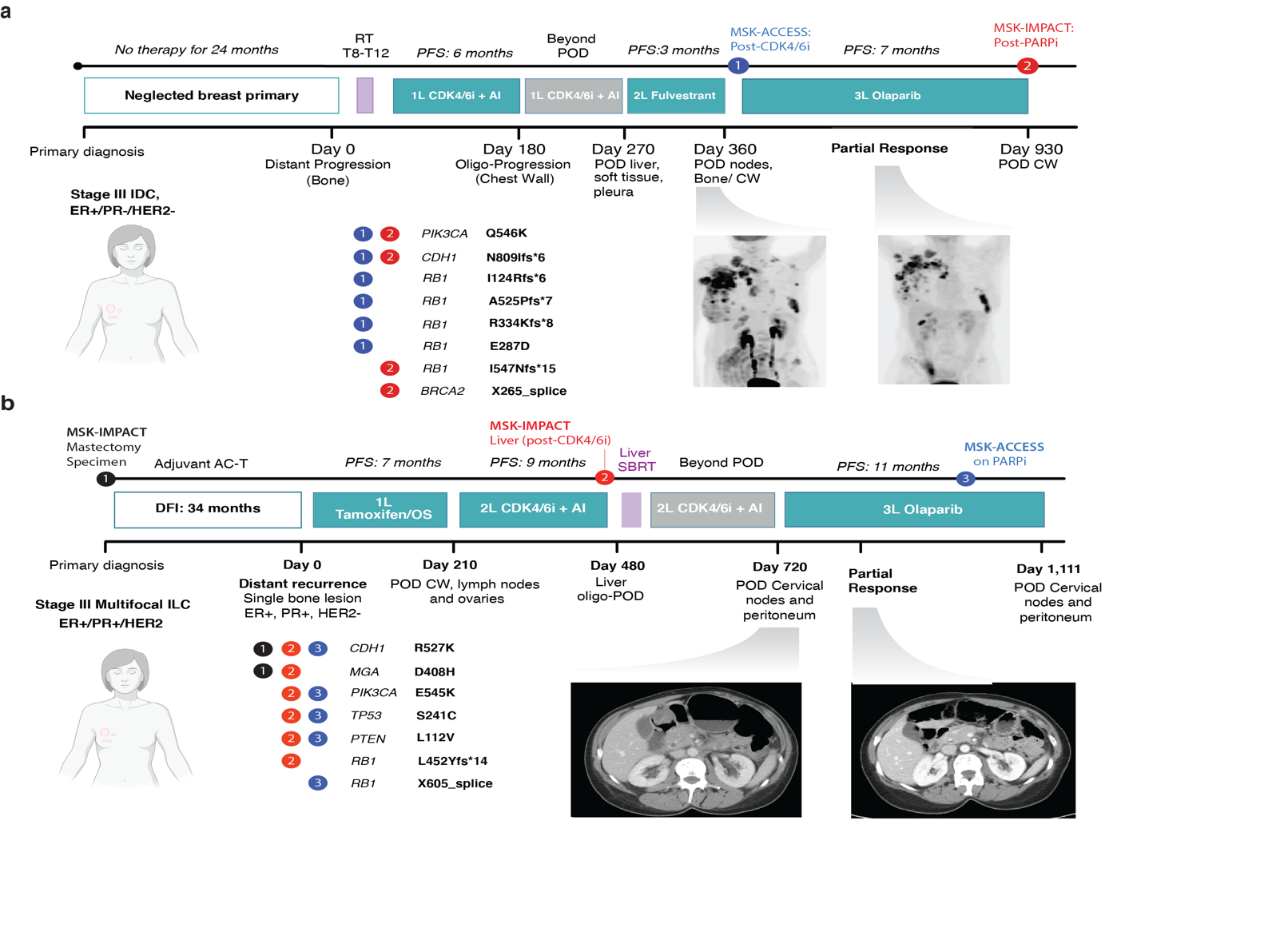

**Fig. S9) Representative cases of short PFS and evidence of convergent *RB1* evolution in g*BRCA2* treated with g*BRCA2* inhibitors. a)** Depicts a patient vignette of locally advanced ER+/PR-/HER2- breast cancer for which patient did not seek treatment for 2 years. She presented with progressive disease in bone and underwent SBRT to T8-T12. She was on CDK4/6i and aromatase inhibitor for 6 months, at which point she experienced oligo-progression in chestwall. These agents were continued for 90 more days, after which progression of disease was noted in parenchyma (liver, pleura, soft musculoskeletal tissue). She was subsequently plaed on fulvestrant, and again experienced progression of disease after 6 months. Post-progression plasma cell free DNA revealed 4 different *RB1* variants. The patient was subsequently treated with olaparib, to which a partial response was noted. She remained on this for 560 days until eventual progression in chest wall. Post progression next generation sequencing of the site of progression revealed a fifth *RB1* loss of functional variant which had not previously been noted. **b)** Depicts a patient vignette for locally advanced lobular carcinoma (ER+/PR+/HER2-) for which adjuvant chemotherapy was pursued. The patient developed distant metastatic recurrence after 34 months, after which she was placed on tamoxifen and ovarian suppression for 7 months. After subsequent progression of disease (chest wall, lymph nodes, ovaries), she was treated with second line CDK4/6i and aromatase inbhitor. She developed oligo progressive disease in the liver. Biopsy revealed a p.L452Yfs*14 variant in *RB1* which was predicted to be loss of function. Subsequent plasma cell-free DNA sequencing revealed an additional splice site variant in *RB1*.

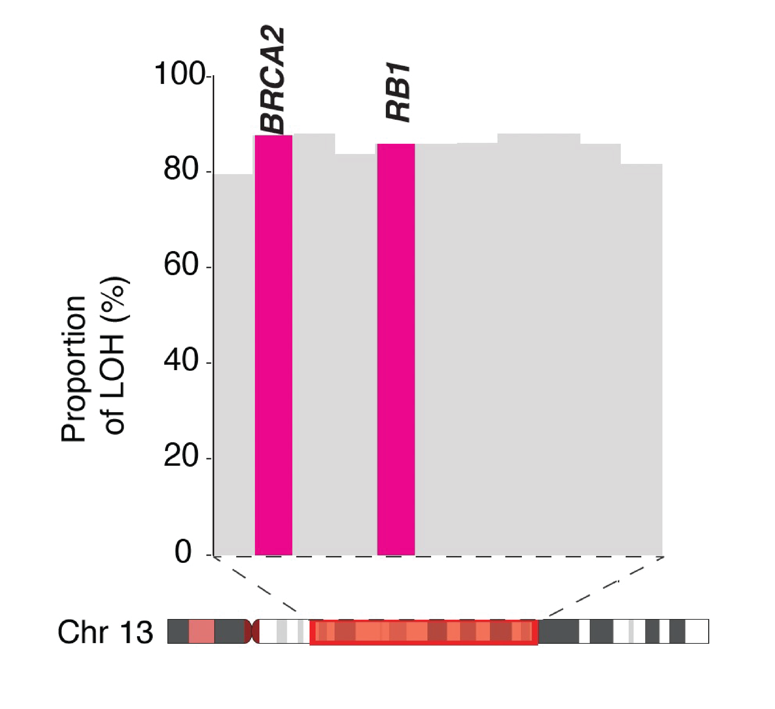

**Fig. S10) Validation of concurrent *RB1* and *BRCA2* LOH in external gBRCA2 WES cohort.** FACETS was performed on whole exome sequencing of breast cancer samples from gBRCA2 Abramson Cancer Center at University of Pennsylvania (n = 24), and Mayo Clinic (n = 22). Of these samples, 38 (82.6%) demonstrated concurrent LOH of *BRCA2* and *RB1*. Two-sided fisher t-test between *RB1* LOH and *BRCA2* LOH was statistically significant (OR = *Inf*, 95% CI 11.12 – *Inf*, p = 3.0 e -6).

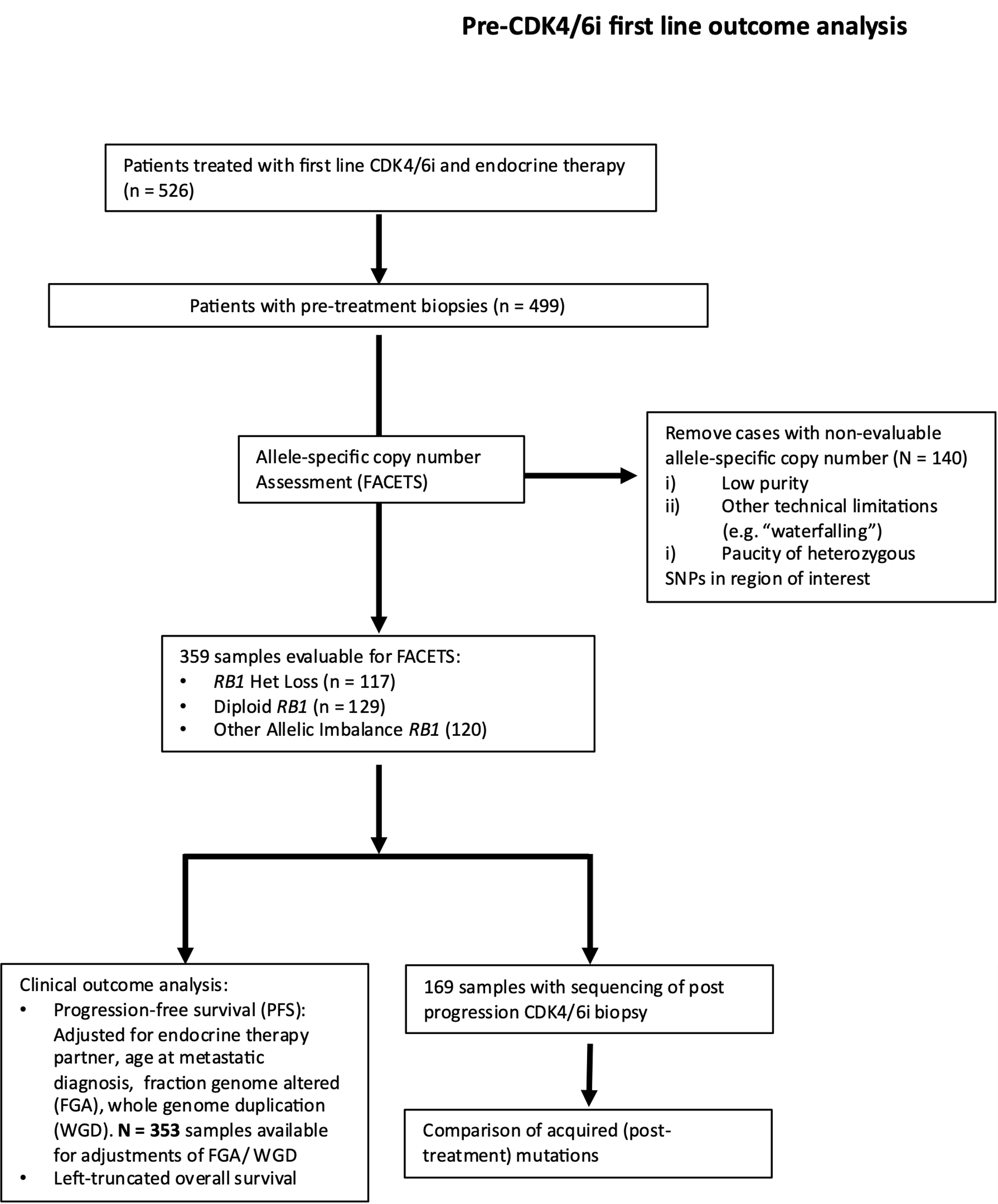

**Fig. S11)** **CONSORT Diagram describing pre treatment allele-specific copy number analysis.** Allele-specific copy number analysis was performed on tumor samples collected prior to initiation of first-line CDK4/6i and endocrine therapy. Samples were excluded (n = 140) for technical reasons including low purity or paucity of heterozygous SNPs in the region of interest. The consort diagram describes the clinical outcomes analysis on the 359 samples remaining, as well as the matched pairs analysis on the 169 samples with corresponding post progression biopsies.
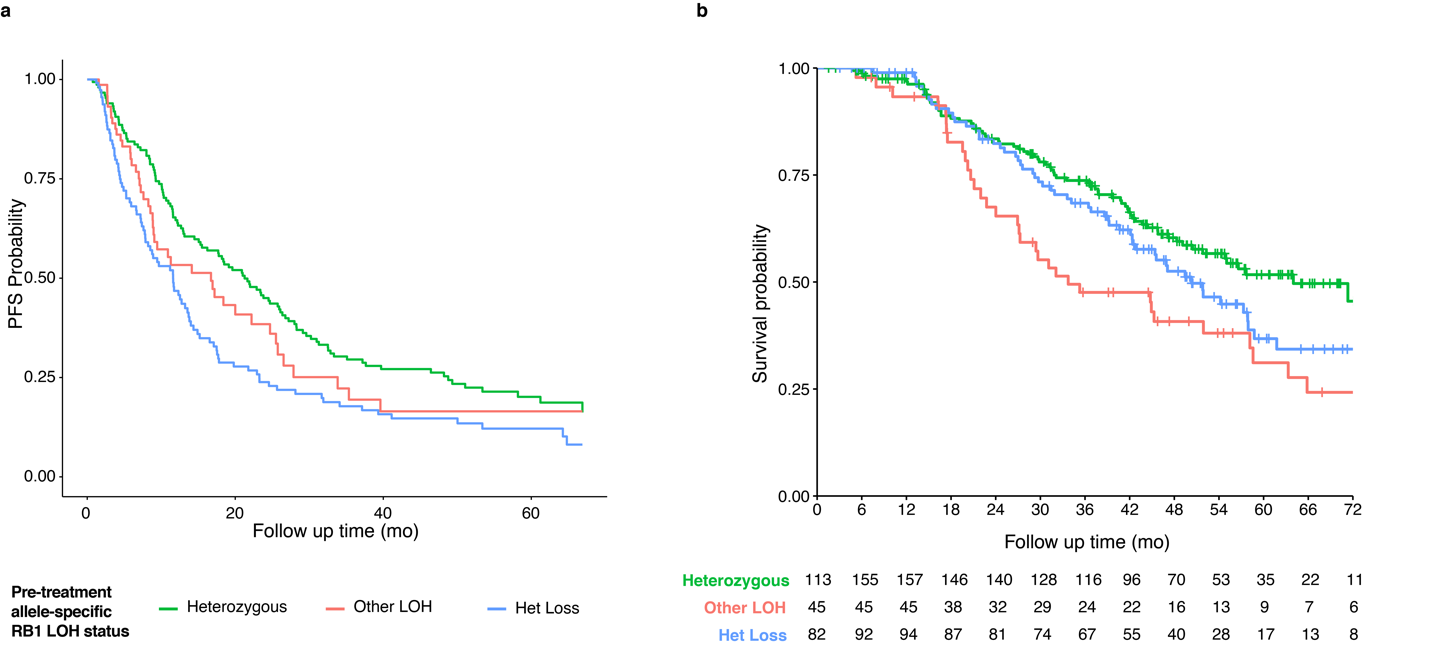

**Fig. S12) Pre-treatment allele-specific copy number analysis. a.** Adjusted Kaplan-Meier curve, accounting for covariates including endocrine therapy partner, age at metastatic diagnosis, fraction genome altered (FGA), whole genome duplication (WGD). After adjusting for relevant genomic and clinical covariates, the group harboring “Other LOH” configurations of RB1 ceases to be associated with PFS (HR 1.35, 95% CI 0.91 – 1.99, p = 0.13). RB1 het loss remains associated with shorter PFS after these adjustments: HR 1.76, 95% CI 1.30 – 2.39, p = 0.00024). **b.** Overall survival curve of patients received first line CDK4/6i and endocrine therapy, based on pre-treatment RB1 allelic configuration. Age at metastatic diagnosis is included as a covariate. “Other LOH” is uniquely associated with a shorter overall survival: HR 1.82, 95% CI 1.19 – 2.79, p = 0.0057. HR of Het Loss: HR 1.32, 95% CI 0.92 – 1.90, p = 0.13).

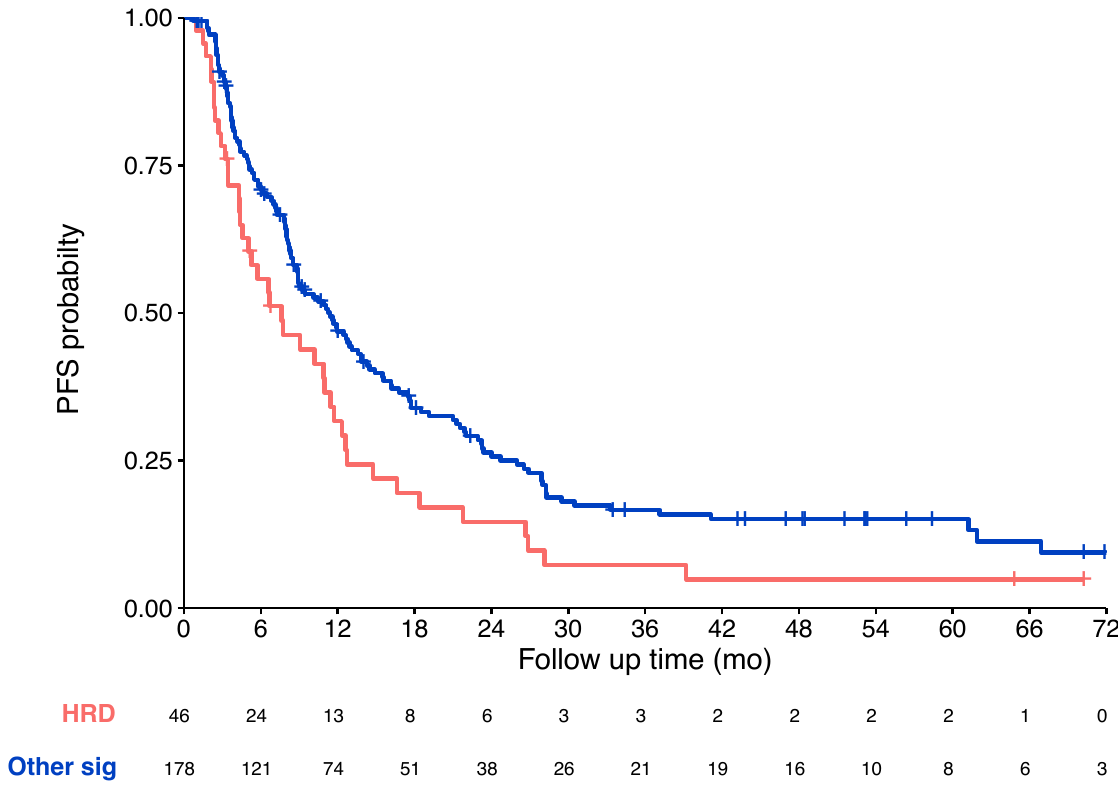

**Fig. S13)** **Implications of HRD signature on CDK4/6i PFS in BRCA2 WT breast cancers**. Signature Multivariate Analysis (SigMA)^48^ was used to derive dominant mutational signature in available cases. Out of 412 cases of pre-CDK4/6i samples (across all treatment signs), 18 g*BRCA2* cases were excluded, and 188 cases were excluded as they did not yield a dominant mutational signature. Compared to non-HRD signature, a dominant HRD signature was associated with decreased progression free survival: HR 1.49, 95% CI 1.04 – 2.12, p = 0.030).
